# Extraction of active RhoGTPases by RhoGDI regulates spatiotemporal patterning of RhoGTPases

**DOI:** 10.1101/720094

**Authors:** Adriana Golding, Ilaria Visco, Peter Bieling, William Bement

**Affiliations:** Graduate Program in Cell and Molecular Biology, University of Wisconsin-Madison, Madison, WI, USA; Department of Systemic Cell Biology, Max Planck Institute of Molecular Physiology, Dortmund, Germany; Laboratory of Cell and Molecular Biology, University of Wisconsin-Madison; Department of Integrated Biology, University of Wisconsin-Madison

## Abstract

The RhoGTPases are characterized as membrane-associated molecular switches cycling between active, GTP-bound and inactive, GDP-bound states. However, 90-95% of RhoGTPases are maintained in a soluble form by RhoGDI, which is generally viewed as a passive shuttle for inactive RhoGTPases. Our current understanding of RhoGTPase:RhoGDI dynamics has been limited by two experimental challenges: direct visualization of the RhoGTPases *in vivo* and reconstitution of the cycle *in vitro*. We developed methods to directly image vertebrate RhoGTPases *in vivo* or on lipid bilayers *in vitro*. Using these tools, we identified pools of active and inactive RhoGTPase associated with the membrane, showed that RhoGDI can actively extract both inactive and active RhoGTPases, and that the extraction of active RhoGTPase contributes to their spatial regulation around wounds. In contrast to the textbook model of the RhoGTPase cycle, these results indicate that RhoGDI actively contributes to spatiotemporal patterning by removing active RhoGTPases from the plasma membrane.

## Introduction

The Rho family GTPases, including Rho, Rac and Cdc42, are essential signaling proteins that mediate morphological changes in cells by directing local cytoskeletal rearrangements (Bishop & Hall, 2000; Kimura et al., 1996). These rearrangements are generally initiated at and confined to specific subcellular regions. For example, a narrow, concentrated zone of Rho activity directs the formation of a ring of actin filaments and myosin-2 at the equatorial cortex that drives cytokinesis (Bement, Benink, and Von Dassow 2005; Yonemura, Hirao-Minakuchi, and Nishimura 2004; Yüce, Piekny, and Glotzer 2005). Similarly, Rho, Rac and Cdc42 are activated near the leading edge of crawling cells in patterns that correspond to local cycles of protrusion, adhesion and retraction (Machacek et al., 2009; Martin et al., 2016). Because tight spatiotemporal regulation of the GTPases is a fundamental feature of these cellular processes, considerable effort has been invested in studying GTPase regulation.

The RhoGTPases are classically characterized as cycling between membrane-associated, active states and soluble, inactive states as a result of interactions with three classes of regulatory proteins: guanine nucleotide exchange factors (GEFs), which activate GTPases by promoting exchange of GDP for GTP (Rossman et al. 2005); GTPase activating proteins (GAPs), which inactivate GTPases by promoting GTP hydrolysis (Moon & Zheng, 2003); and guanine nucleotide dissociation inhibitor (GDI), which solubilizes GTPases to generate a large reservoir of heterodimeric GTPase:GDI complexes in the cytoplasm (Garcia-Mata, Boulter, & Burridge, 2011). In the canonical model of GTPase regulation, GTPase cycling is thought to proceed as follows: a GTPase is activated by a GEF at the plasma membrane following its release from GDI, is subsequently inactivated by a GAP, and is then returned to the soluble pool by GDI. Thus, in the canonical model, the lifetime of GTPase activity at the plasma membrane is thought to be controlled entirely by GEFs and GAPs, with GDI essentially serving as a passive shuttle that interacts exclusively with inactive GTPases.

A limitation of this traditional view is that the function and the biochemical activities of RhoGDI, in contrast to GEFs and GAPs, are not well understood. Presently, no consensus exists as to i) whether GDI actively extracts GTPases from membranes or merely solubilizes them by sequestration (Johnson, Erickson, and Cerione 2009; Zhang et al. 2014), ii) whether it interacts with GTPases in a nucleotide-specific manner (Nomanbhoy & Cerione, 1996; Tnimov et al., 2012), or iii) how its activity is coordinated with GEFs or GAPs (Garcia-Mata et al., 2011). The notion that GDI works as a passive shuttle rests largely on two findings. First, when GTPase:GDI complexes are purified from cell lysates, the great majority of GTPase within the complex is in the inactive, GDP-bound form (Abo, Webb, Grogan, & Segal, 1994), as expected if the GDI solubilizes GTPases after inactivation by a GAP. Second, binding of GTPases by GDI strongly suppresses GTP hydrolysis (Hart et al., 1992), indicating that hydrolysis must precede the extraction from the membrane. However, other results raise a serious, albeit contentious, challenge to this idea: multiple studies indicate that GDI binds both inactive and active GTPase with relatively high affinity *in vitro* (Hancock & Hall, 1993; Hart et al., 1992; Nomanbhoy & Cerione, 1996; Tnimov et al., 2012), implying that GDI may have the potential to interact with active as well as inactive GTPase *in vivo*. As such, GDI might have the potential to exert a more direct role in the regulation of GTPase activity than currently appreciated.

Understanding the pattern forming ability of RhoGTPases requires the mechanistic dissection of the interactions between GTPases and GDI. Unfortunately, our ability to study GTPase:GDI dynamics has been hampered due to two major experimental limitations: first, visualization of the GTPases in living cells is limited by the fact that labeling with fluorescent protein on the amino terminus impairs GTPase regulation and function, while carboxy-terminal labeling prevents GTPase prenylation (Howell et al., 2012; Yonemura et al., 2004). In the absence of direct visualization, GTPase dynamics must be inferred from activity probes. Second, with a few important exceptions (Johnson et al., 2009; Nomanbhoy, Erickson, & Cerione, 1999), *in vitro* studies of GTPase:GDI dynamics have either utilized unprenylated GTPases, omitted membranes, or both. Additionally, nearly all reconstitution experimentation focused on the effect of GDI on the distribution between membrane-associated and soluble forms of GTPases at thermodynamic equilibrium (Zhang et al., 2014). Thus, we do not currently understand how GDI affects the transitions between membrane and soluble GTPase states kinetically, especially under conditions which mimic the cellular environment which is far from equilibrium due to the constant dissipation of energy.

To overcome these limitations, we developed two distinct methods to directly visualize vertebrate RhoGTPases on membranes *in vivo* and or on supported lipid bilayers *in vitro*. Using these tools, we identify co-existing pools of active and inactive GTPases associated with the plasma membrane. We also demonstrate that GDI can actively extract GTPases from the membrane and, unexpectedly, that GDI can extract both inactive and active GTPases. Finally, we show that the extraction of active GTPase also occurs *in vivo* and that this contributes to the spatial regulation of GTPase activity. Collectively, these data indicate that the textbook model of the GTPase cycling must be reassessed because GDI can directly mediate the spatiotemporal regulation of GTPase activity.

## Results

### Visualization of RhoGTPases around cell wounds

Traditional fusion of fluorescent protein with the amino- or carboxyl-termini of the RhoGTPases impairs GTPase localization and function (SuppFig1; Yonemura, Hirao-Minakuchi, and Nishimura 2004). To overcome this problem, we first adapted an approach described by Bendezú et al. (2015) for labeling of yeast Cdc42. Specifically, we inserted green fluorescent protein into a solvent-exposed external loop of the *Xenopus* GTPases (see Methods). To test the internally-tagged (IT) GTPases *in vivo,* we exploited the cell wound repair model in *Xenopus laevis* oocytes where wounding elicits a robust accumulation of active Rho and Cdc42 in discrete, concentric zones at the cortex (Fig1A; Benink & Bement, 2005). IT-GTPases were co-expressed with wild-type (WT) GDI to avoid GTPase aggregation (Boulter et al., 2010). Both IT-Rho and IT-Cdc42 were recruited to concentric rings around the wound (Fig1B,C). Comparison of IT-Rho to a Rho activity reporter (mRFP-2xrGBD; Davenport et al. 2016) revealed that IT-Rho spatially overlapped with the Rho activity zone. Comparison of IT-Cdc42 to a Cdc42 activity reporter (mRFP-wGBD; Benink and Bement 2005) revealed that IT-Cdc42 localized throughout the active Cdc42 zone, but extended slightly beyond it towards the wound center (Fig1B,C; see also below). We also tested the behavior of IT-Rac and found that it concentrated around wounds in the same region as IT-Cdc42 as expected from previous experiments (SuppFig2; Abreu-Blanco, Verboon, & Parkhurst, 2014; Benink & Bement, 2001). These results indicate that the IT and Cy3-tagged GTPase variants can interact with diverse regulators required to achieve their normal enrichment at wounds.

**Figure 1:**
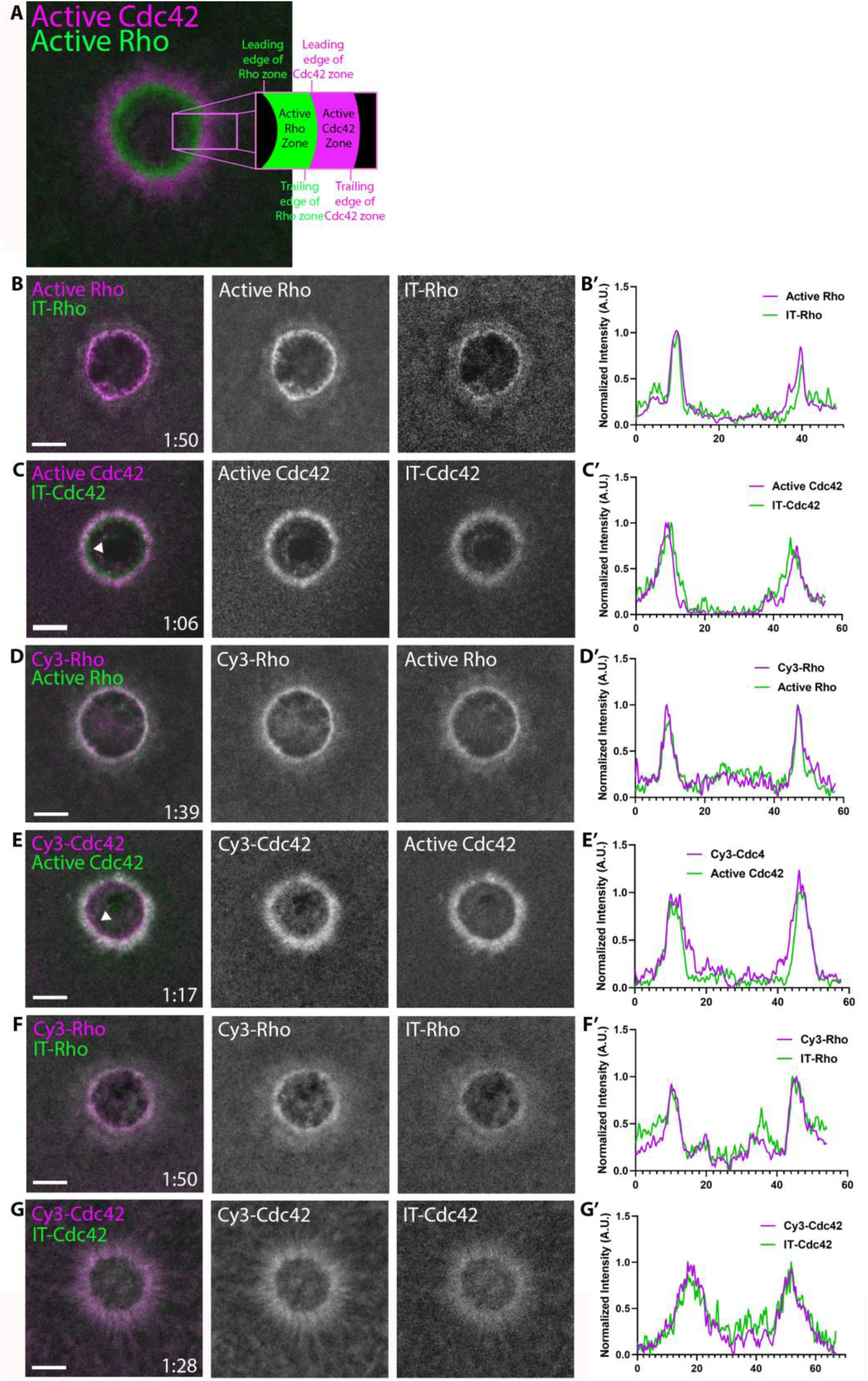
Direct visualization of Rho and Cdc42 during cell wound repair. A) Left: image of active Cdc42 (magenta) and active Rho (green) around a single-cell wound; right: schematic diagram indicating zone regions; B) Wound in oocyte microinjected with rGBD (magenta) and IT-Rho (green); B’) Line scan of normalized fluorescence intensity from (B); C) As in B but with wGBD (magenta) and IT-Cdc42 (green); D,D’) As in B but with Cy3-Rho (magenta) and rGBD (green); E,E’) As in B but with Cy3-Cdc42 (magenta) and wGBD (green); F,F’) As in B but with Cy3-Rho (magenta) and IT-Rho (green); G,G’) As in B but with Cy3-Cdc42 (magenta) and IT-Cdc42 (green) and line scan. Scale bar 10μm, time min:sec.

As an alternative approach, and as a means to obtain labeled RhoGTPases that could be used both *in vivo* and *in vitro*, purified recombinant Rho and Cdc42 were prenylated and coupled to Cy3 via a short N-terminal peptide via sortase-mediated ligation (see Methods). Microinjected Cy3-Rho and Cy3-Cdc42 localized to wounds (Fig1D,E), in a manner indistinguishable from their IT counterparts expressed in the oocyte (Fig1F,G). As observed with IT-Rho and IT-Cdc42, Cy3-Rho completely overlapped with the zone of Rho activity while Cy3-Cdc42 localized throughout and slightly interior of the active Cdc42 zone.

### Visualization of RhoGTPases in other cellular contexts

To further test the behavior of the IT- and Cy3-labeled RhoGTPases, we sought to determine if they localize to the plasma membrane in other cellular processes. This is important because these processes likely depend on different regulators from those that operate during cell wound repair. IT-Rho and Cy3-Rho localized to the cytokinetic apparatus and epithelial junctions in *Xenopus* embryos, consistent with previous results obtained with a Rho activity reporter (Fig2A-C; Bement, Benink, & Von Dassow 2005). Similarly, IT-Cdc42 localized to exocytosing cortical granules (Fig2D), consistent with previous results obtained with a Cdc42 activity reporter (Yu and Bement 2007). IT-Cdc42 was also recruited to cell-cell junctions and enriched there upon wounding (Fig2E), a behavior previously revealed using a Cdc42 activity reporter (Clark et al., 2009).

**Figure 2:**
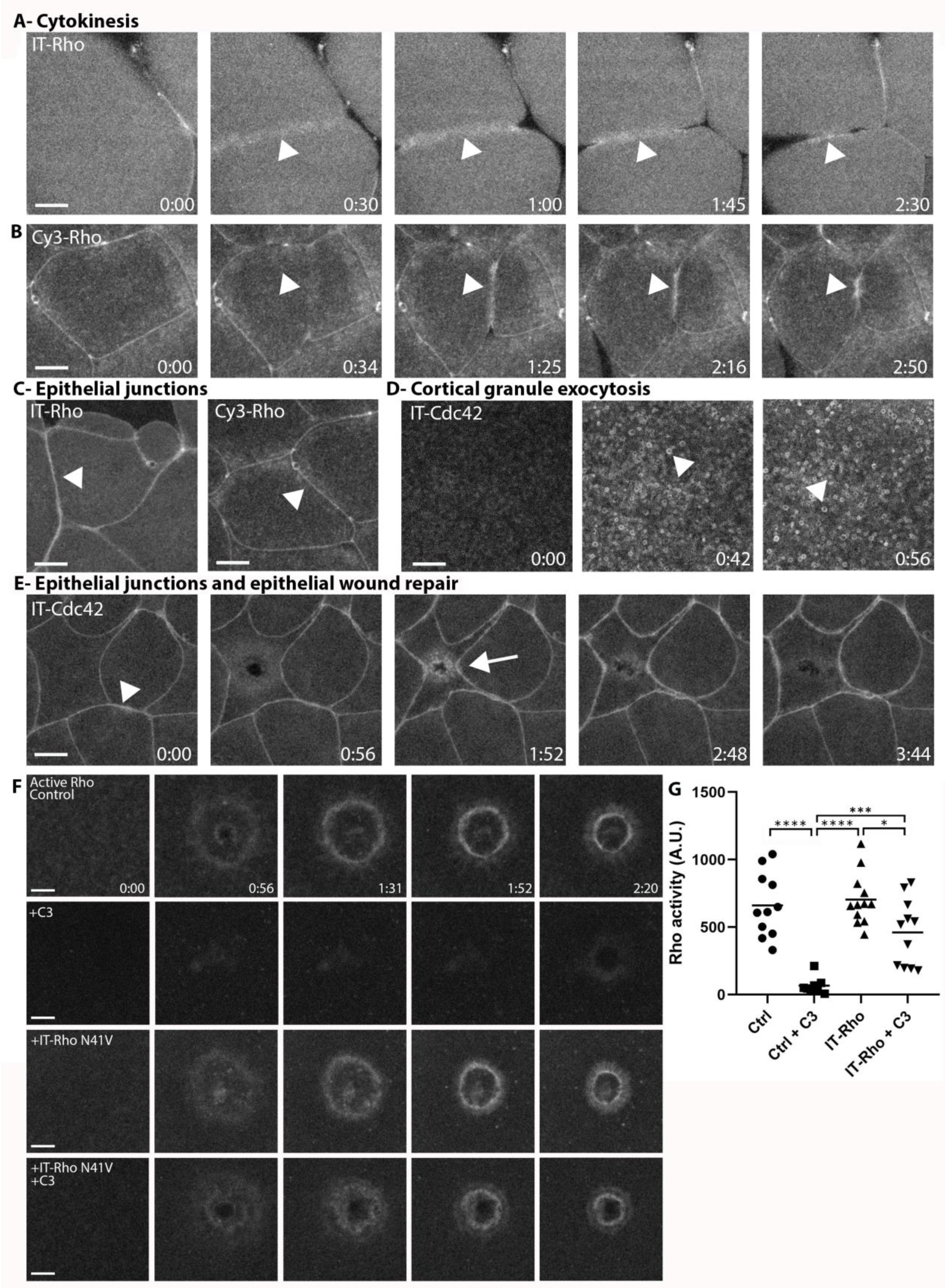
Directly-labeled Rho and Cdc42 during cytokinesis, cortical granule exocytosis, epithelial wound repair and at junctions. A) Cytokinesis in *X. laevis* embryo microinjected with IT-Rho; IT-Rho accumulates at nascent cleavage furrow (arrowhead); B) Cytokinesis in *X. laevis* embryo microinjected with Cy3-Rho; Cy3-Rho accumulates at nascent cleavage furrow (arrowhead); C) *X. laevis* embryo microinjected with IT-Rho (left) and Cy3-Rho (right); both are enriched at cell-cell junctions; D) Meiotically mature *Xenopus* egg microinjected with IT-Cdc42; IT-Cdc42 is recruited to exocytosing cortical granules (arrowheads) following egg activation (0:42); E) *X. laevis* embryo microinjected with IT-Cdc42; IT-Cdc42 concentrates at cell-cell junctions (arrowhead; 0:00) and, following damage, is recruited to the wound and becomes enriched at junctions (arrow); F) C3-insensitive IT-Rho rescues Rho activation in presence of C3. Control: cell microinjected with rGBD shows normal Rho activity accumulation and wound closure; C3: cell microinjected with rGBD fails to activate Rho in presence of C3; IT-Rho-N41V: cell microinjected with rGBD and C3-insensitive IT-Rho normally activates Rho; IT-Rho-N41V+C3: cell microinjected with rGBD and C3-insensitive IT-Rho rescues Rho activity in presence of C3.Scale bar 10μm, time min:sec. G) Quantification of Rho activity, corrected for background (n=8-12). One-way ANOVA with Tukey post-test statistical analysis. *p<0.05, ***p<0.001, ****p<0.0001.

Next, we wanted to determine whether IT-RhoGTPases can functionally substitute for the endogenous GTPases. The *Xenopus laevis* oocyte system is not conducive to traditional knockdown approaches due to its large stores of maternal protein and relatively slow protein turnover. Therefore, we employed C3-exotransferase, a Rho-specific toxin, to inhibit endogenous Rho activity, and expressed an IT-Rho in which the C3 ribosylation site (N41) is mutated to a residue that cannot be ribosylated (N41V; Sekine, Fujiwara, and Narumiyas 1989). In control oocytes expressing the probe for active Rho, Rho activity around wounds was suppressed by C3 (Fig 2F,G). In contrast, cells expressing IT-Rho-N41V generated a spatially defined zone of Rho activity around the wound that closed over similar timescales compared to the control. Collectively, the above results indicate that both IT- and Cy3-labeled GTPases are faithful reporters of the distribution of GTPases and show that IT-Rho can functionally substitute for its endogenous counterpart.

### Pools of active and inactive RhoGTPase accumulate in the plasma membrane

The observation that Cdc42 extended slightly beyond its zone of activity suggests that there may be a pool of inactive, membrane-bound Cdc42 at this location. This notion is consistent with the previous observation that Abr, a Cdc42 GAP thought to regulate Cdc42 activity at wounds, also localizes interior to the Cdc42 zone (Vaughan et al. 2011). To understand the relationship between activity and membrane-association of Cdc42, we overexpressed Abr, a manipulation previously shown to decrease Cdc42 activity around wounds (Vaughan et al., 2011). Remarkably, this resulted in a dose-dependent loss of active Cdc42 at wounds while having far less effect on Cy3-Cdc42 (Fig3A,B). These results further demonstrate that the IT- and Cy3-GTPases are functional. More importantly, they demonstrate that substantial pools of both active and inactive GTPases can be dynamically maintained at the plasma membrane.

**Figure 3:**
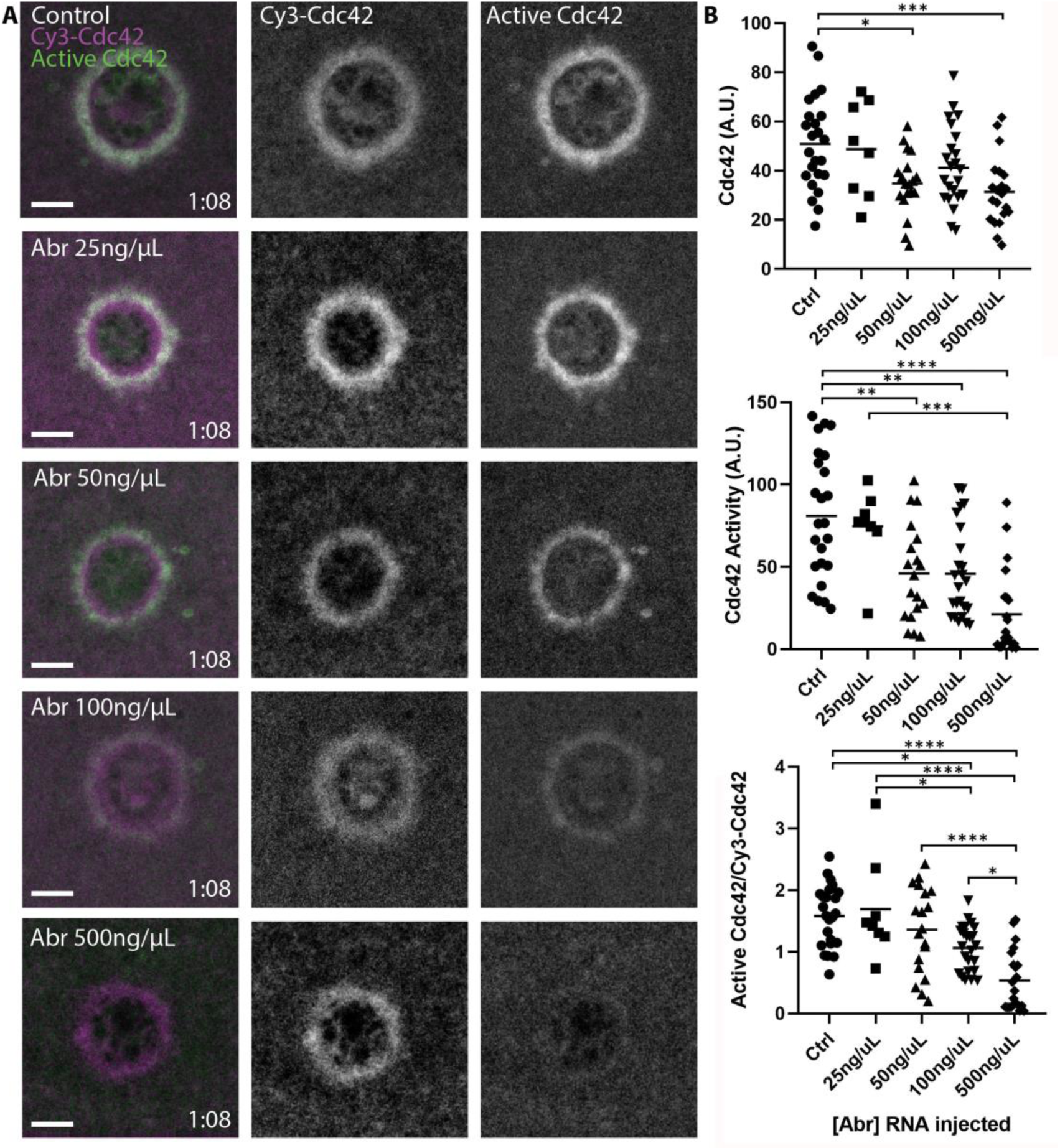
Pools of inactive and active Cdc42 at the plasma membrane. A) Oocytes microinjected with wGBD (green), Cy3-Cdc42 (magenta) and indicated concentrations of mRNA encoding the Cdc42-GAP Abr. Scale bar 10μm, time min:sec.; B) Quantification of Cy3-Cdc42, active Cdc42 and ratio of active Cdc42 to Cy3-Cdc42 for each condition. n=8-24. One-way ANOVA with Tukey post-test statistical analysis. *p<0.05, **p<0.01, ***p<0.001, ***p<0.0001.

### RhoGDI is recruited to concentrated areas of RhoGTPase activity

Efforts to visualize RhoGDI at the plasma membrane have generally failed (Ngo et al., 2017), likely because GDI only transiently interacts with GTPases upon release into or extraction from the membrane. However, we reasoned it might be possible to detect GDI at wound sites due to the high local concentration of Rho and Cdc42. Indeed, we found that 3xGFP-GDI is enriched at wounds and overlaps with both the Rho and Cdc42 zones (Fig4A). To confirm that endogenous GDI also localizes to wounds, antibodies were raised against *X. laevis* GDI (SuppFig3A) and used to immunolabel wounded oocytes. Consistent with the results obtained with 3XGFP-GDI, endogenous GDI accumulated at wounds (Fig4B). These results demonstrate that GDI accumulation occurs at discrete regions of the plasma membrane that are enriched with its GTPase clients.

**Figure 4:**
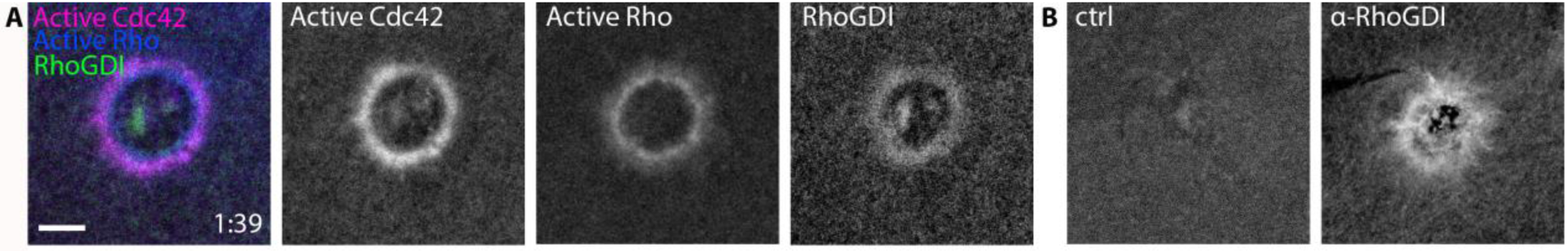
RhoGDI is recruited to single-cell wounds enriched in Rho and Cdc42 activity. A) Oocytes microinjected with wGBD (magenta), rGBD (blue) and GDI (green); B) Wounded oocytes fixed and stained with anti-*X. laevis* RhoGDI. Scale bar 10μm, time min:sec.

### RhoGDI differentially regulates Rho and Cdc42

The localization of RhoGDI around wounds suggests that it might play an active role in delivery to or extraction of RhoGTPases from the membrane and thus their spatiotemporal patterning. As an initial test of this possibility, we overexpressed GDI via mRNA microinjection. This manipulation potently suppressed both Rho and Cdc42 activity, as well as Cy3-Rho and Cy3-Cdc42 localization, suggesting that GDI exerts its effects via extraction of the GTPases (Fig5A). To obtain a more quantitative understanding of the relationship between GDI and GTPase activity, we microinjected purified GDI (SuppFig3B) at increasing concentrations prior to wounding. High concentrations of microinjected GDI suppressed both Rho and Cdc42 activity at wounds (Fig5B,C), consistent with the results obtained from GDI via mRNA-mediated overexpression. However, more modest increases revealed differential effects on Rho and Cdc42. Specifically, increases between 18-190% in GDI levels (based on proteomic data from Wühr et al. 2014) resulted in a greater reduction of Cdc42 activity compared to Rho (Fig5C). We found the same to be true for bovine GDI (SuppFig4). These results show that GDI differentially impacts Rho and Cdc42 activity *in vivo* and that this effect does not require gross overexpression.

**Figure 5:**
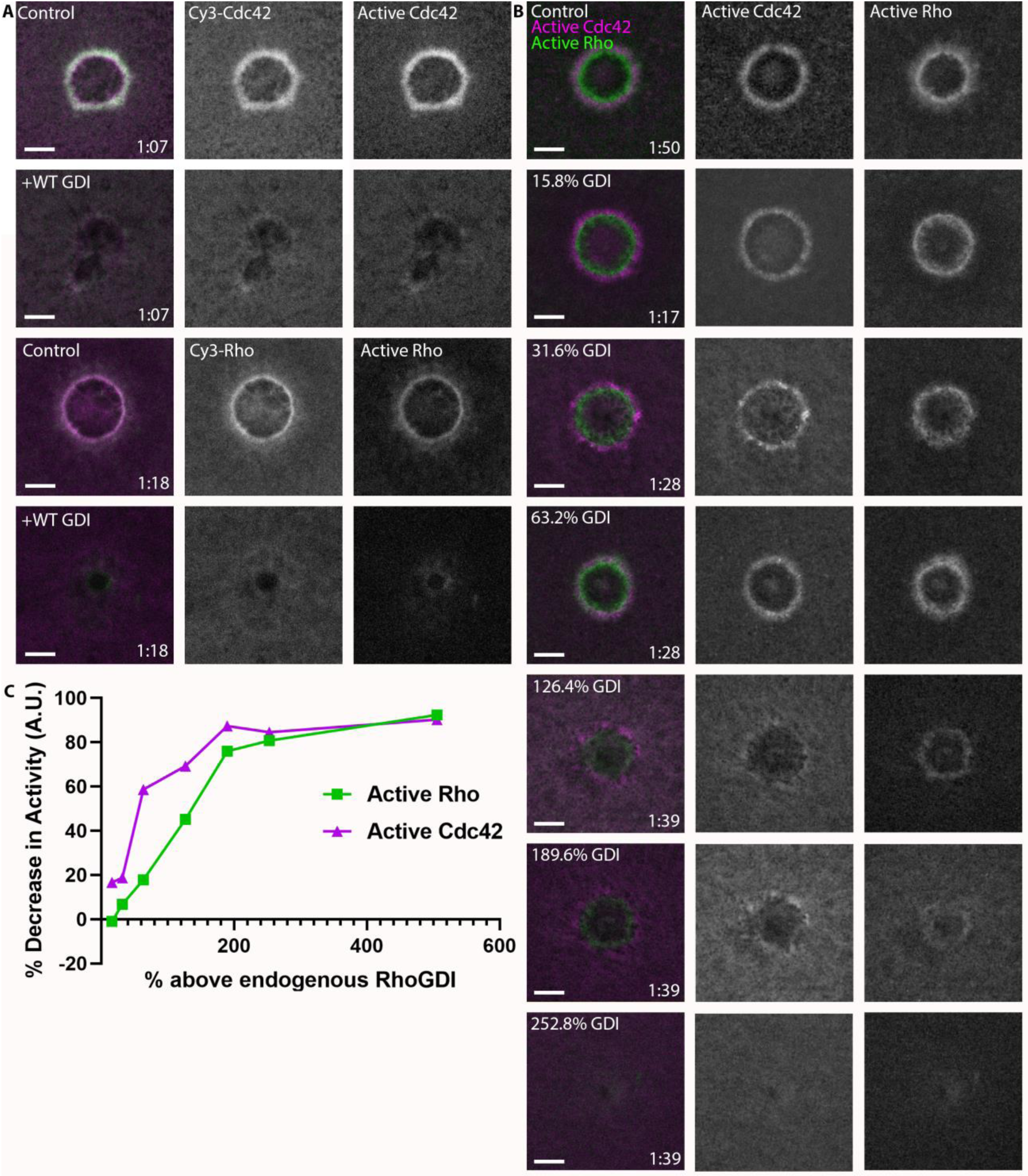
RhoGDI overexpression differentially regulates Rho and Cdc42 activity. A) Top 2 rows: oocytes microinjected with Cy3-Cdc42 (magenta) and wGBD (green) alone or with RhoGDI. Bottom 2 rows: Oocytes microinjected with Cy3-Rho (magenta) and rGBD (green) alone or with GDI; B) Oocytes microinjected with wGBD (magenta), rGBD (green) and increasing concentrations of RhoGDI protein; C) Standard curve of decrease in Rho and Cdc42 activity with increasing concentrations of RhoGDI (n=10-23 for each concentration). Scale bar 10μm, time min:sec.

### RhoGDI extracts active and inactive RhoGTPase in vitro

To directly probe the mechanism by which RhoGDI inhibits Rho and Cdc42 activity *in vivo*, we established a real-time GTPase dissociation assay on supported lipid-bilayers (SLBs) (Fig6A; SuppFig5; see Methods). The addition of Cy3-Cdc42 to SLBs resulted in their membrane binding as detected by total internal reflection microscopy (TIRF), which was dependent on its C-terminal prenyl moiety, as expected (Fig6B). We then studied the time course of Cdc42 release from SLBs under buffer flow which continuously flushed out unbound proteins from solution. Spontaneous release of inactive, GDP-bound Cdc42 from the membrane was rather slow (t1/2=37.24±3.05 sec), however the addition of excess GDI lead to a dramatic acceleration of dissociation (ca.20-fold; Fig6C; SuppFig3C). To determine whether this was the result of either simple sequestration in solution or, alternatively, active catalytic extraction of Cdc42 from membranes, we performed assays in the presence of an alternative solubilizer (RabGGTase beta) that sequesters the GTPase prenyl moiety (Fig6C). Sequestration alone only marginally affected the rate of dissociation, demonstrating that GDI actively extracts GTPases from membranes (Fig6D,E).

**Figure 6:**
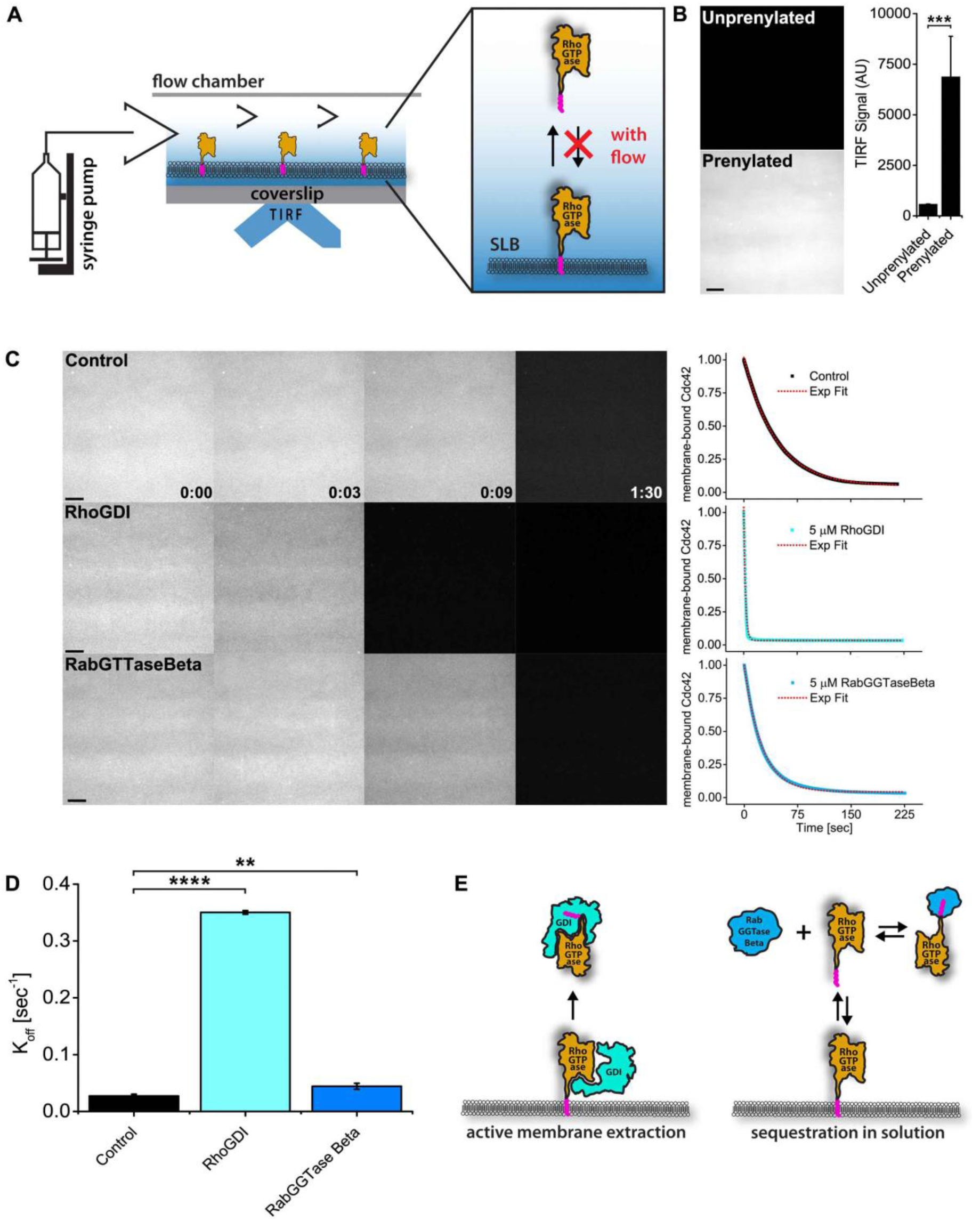
RhoGDI actively extracts RhoGTPases from membranes *in vitro*. A) Experimental setup of *in vitro* experiments: prenylated RhoGTPases were reconstituted on supported lipid bilayers (SLBs) in flow chambers and imaged by TIRF. Wash off experiments were designed to avoid RhoGTPase rebinding to membranes and performed controlling the flow rate via a syringe pump; B) TIRF imaging allows for selective imaging of RhoGTPases at the membrane. Prenylated and unprenylated Cdc42 were imaged in the same conditions and TIRF signal at membranes was quantified (n=4 for each condition); C) Wash off experiments: prenylated Cdc42 reconstituted on SLBs were washed with imaging buffer only (control), in presence of 5μM RhoGDI or RabGTTase Beta. Time lapse images at selected time points and quantification of the full experiments are shown. Decay curves were fitted with a monoexponential function; D) Comparison of the K_off_ values obtained by fitting the decay curves (n=3 for control and RabGTTase Beta, n=2 for RhoGDI); E) Schematic representation of the proposed mode of action of the two RhoGTPases solubilizers. RabGGTase Beta sequesters RhoGTPases in solution, whereas RhoGDI actively extracts RhoGTPases from the membranes. Scale bar 10μm. Unpaired student’s t-test, 2-tailed distribution, equal variance statistical analysis. **p<0.01, ***p<0.001, ****p<0.0001.

To characterize membrane extraction more quantitatively, we carried out experiments over a wide range of RhoGDI concentrations using either inactive (GDP-bound) or active (GTP-bound, constitutively active) forms of Cdc42 or Rho. We observed that WT RhoGTPases hydrolyze even GTP analogs such as GTPγS over the long time period (hours) required for performing a full titration in our SLB assay (data not shown), which lead to them accumulating in the inactive, GDP-bound form during the course of the experiment. We therefore turned to two constitutively active GTPase variants -Q61L and G12V- and found that the former was the most suitable for our in vitro assay because of its extremely low rate of spontaneous GTP hydrolysis. Nonetheless, G12V produced qualitatively similar results in this assay (see SuppFig5 and Supplemental Discussion). Remarkably, GDI was able to extract both inactive (GDP-bound) and active (GTP-bound, Q61L/Q63L) Cdc42 and RhoA in a concentration-dependent manner (Fig7A,B). Although GDI extracted GDP-bound GTPase more efficiently than GTP-bound GTPases (3-5 fold difference), it was still able to effectively facilitate the dissociation of the latter (Figure 7C-F). The affinities of GDI for the active and inactive GTPases on membranes, determined by hyperbolic fits to the extraction rates, were surprisingly similar (Cdc42:GDP 3.41±0.56µM, Cdc42Q61L:GTP 14.40±2.50µM, RhoA:GDP 5.77±0.87µM, RhoAQ63L:GTP 19.58±2.45µM). On the other hand, the maximal rates of extraction were not, indicating that the rate-limiting step of membrane extraction depends on the activity state of GTPases. To investigate whether extraction of active and inactive GTPases is a conserved ability among GDI proteins, we also studied mammalian GDI. Similar to its *Xenopus* ortholog, bovine GDI1 was able to extract both GDP- as well as GTP-bound Cdc42 and RhoA (SuppFig6). These data clearly demonstrate that GDIs can catalytically extract both inactive and active RhoGTPase from membranes *in vitro*.

**Figure 7:**
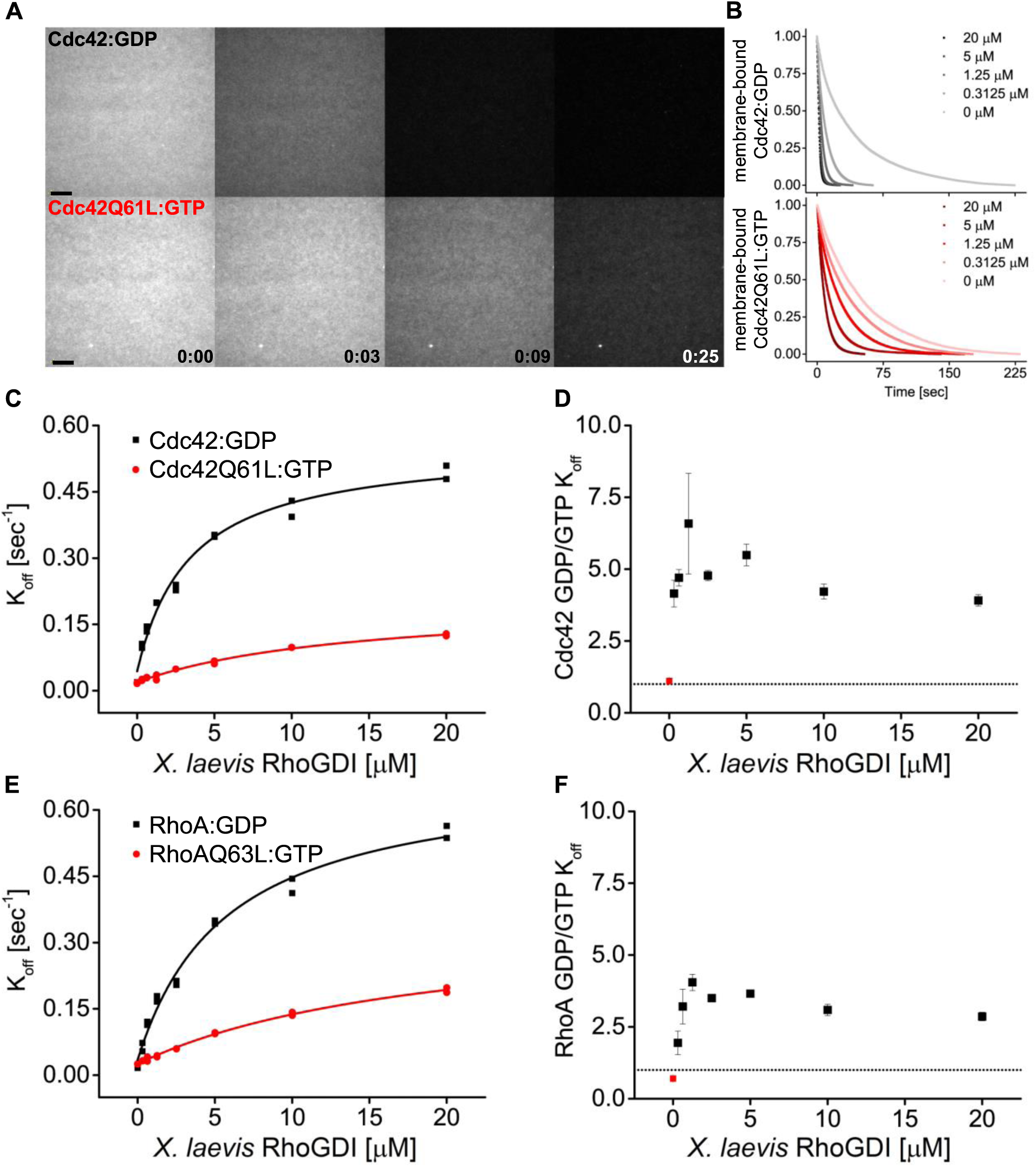
RhoGDI actively extracts both inactive and active RhoGTPases from membranes *in vitro*. A) Wash off experiments: prenylated Cdc42 in both inactive (Cdc42:GDP) and constitutively active (Cdc42Q61L:GTP) states were reconstituted on SLBs and washed in presence of 5 μM RhoGDI. Time lapse images at selected time points are shown; B) Quantification of wash off experiments in which the concentration of RhoGDI was titrated between 0 and 20μM; C) K_off_ values obtained for inactive and constitutively active Cdc42Q61L fitting the decay curves with a monoexponential decay function are plotted against RhoGDI concentration. Extraction rates were fitted with a hyperbolic function; D) Ratio of K_off_ obtained for inactive and constitutively active Cdc42Q61L at the same RhoGDI concentration; F) Same as in C-D for inactive (RhoA:GDP) and constitutively active (RhoAQ63L:GTP) RhoA. Scale bar 10μm.

### Identification of an extraction-deficient RhoGDI

The canonical RhoGTPase cycle assumes that GDI does not extract GTPase without its prior inactivation by a GAP (*reviewed by* Garcia-Mata, Boulter, & Burridge 2011). However, the above results suggest that GDI might directly attenuate GTPase activity at the plasma membrane via extraction of active GTPase. If this hypothesis is correct, then expression of an extraction-deficient GDI would influence the spatiotemporal patterning of GTPase activity. We therefore sought to generate an extraction-deficient GDI that could still initiate contact with GTPases but fail to extract them from the plasma membrane. Mutants were screened by quantifying their recruitment to wounds relative to WT GDI, based on the rationale that mutants capable of binding but not extracting should remain at the membrane longer and thus accumulate at wounds more than WT GDI.

Using this screen, we first tested RhoGDI mutants that were previously reported to be deficient in extraction (Dransart, Morin, Cherfils, & Olofsson, 2005; Ueyama et al., 2013). None of these extraction-deficient mutants were recruited to wounds more strongly than WT GDI, suggesting that they were impaired in binding to GTPases at wounds rather than extraction (SuppFig7). We therefore designed three novel mutants: (1) the isolated regulatory arm of GDI that initiates contact with the GTPase but lacks a binding pocket for the hydrophobic prenyl group (Δ51-199) and (2,3) mutation of residues E158/9 previously hypothesized to be responsible for GTPase extraction (Hoffman, Nassar, & Cerione, 2000). GDI mutants Δ51-199, E158/9A and E158/9Q were halo-tagged, expressed in oocytes and their recruitment to wounds was quantified relative to WT GDI. GDI Δ51-199 showed minimal recruitment to wounds, however both GDI E158/9A and E158/9Q showed a significant increase in recruitment to wounds relative to WT GDI, with GDI E158/9Q (GDI-QQ) having the greatest increase (Fig8A,B).

**Figure 8:**
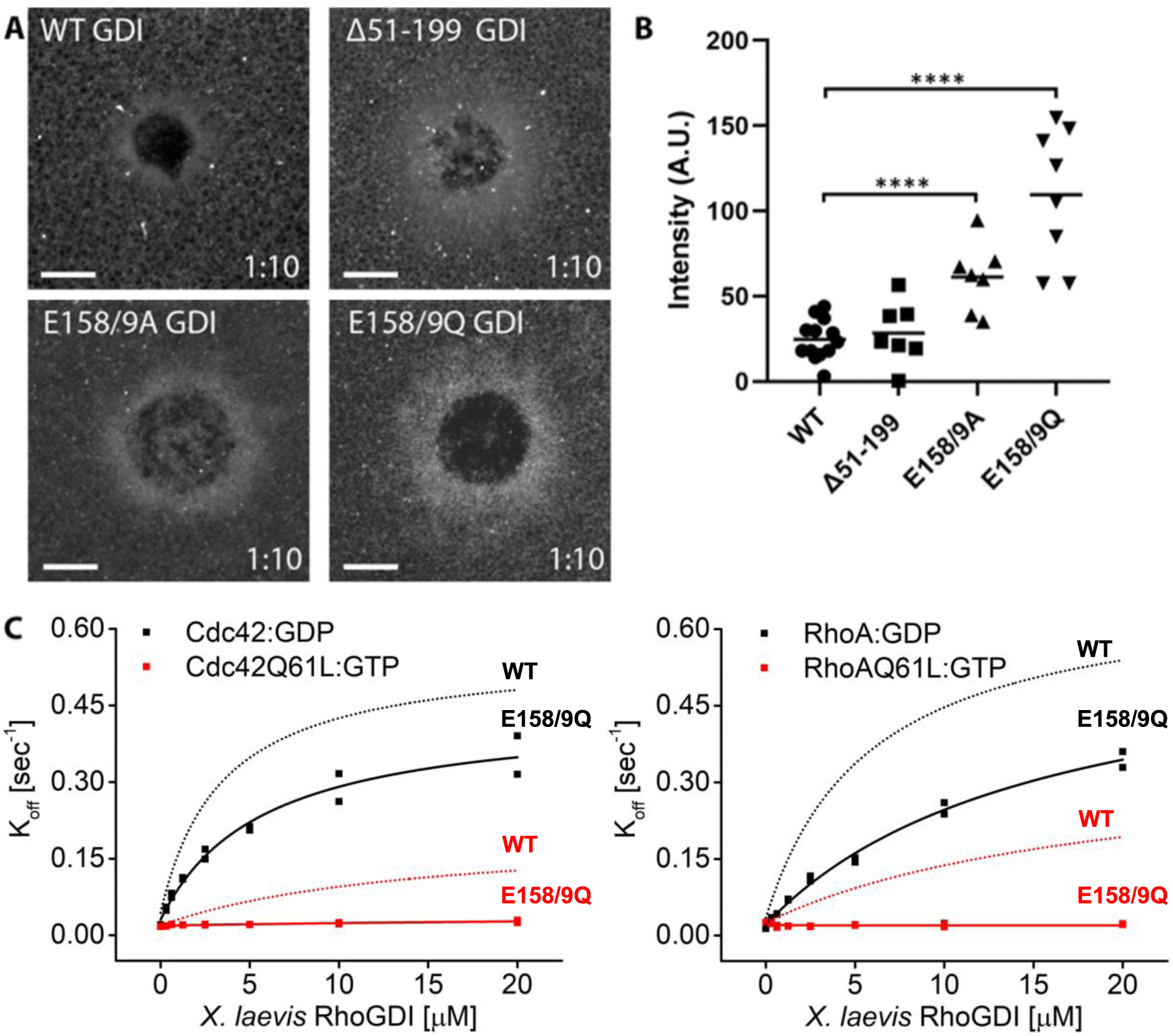
Identification of mutant RhoGDI deficient in extraction of active RhoGTPase. A) Oocytes microinjected with halo-tagged WT, Δ51-199, E158/9A and E158/9Q GDI mutants. Scale bar 10μm, time min:sec; B) Quantification of RhoGDI intensity at wounds (n=7-13). Unpaired student’s t-test, 2-tailed distribution, unequal variance statistical analysis to WT. ****p<0.0001. C) Comparison of K_off_ values obtained for inactive (Cdc42:GDP, RhoA:GDP) and constitutively-active (Cdc42Q61L:GTP, Cdc42Q63L:GTP) Cdc42 and RhoA from wash off experiments in presence of either WT (black) or E158/9Q (QQ) (red) RhoGDI. Extraction rates were fitted with a hyperbolic function.

To directly test whether RhoGDI E158/9Q (QQ) is deficient in extraction, its functional capabilities were tested *in vitro* in the SLB assay. WT GDI was able to extract both GDP-bound and GTP-bound Cdc42 Q61L from the supported lipid bilayers (Fig8C). In contrast, GDI-QQ retained most of its ability (less than two-fold reduction) to extract inactive Cdc42 but was completely deficient in extracting active Cdc42 (Fig8C). Quantitatively similar results were obtained using Cdc42 G12V (data not shown). The same was found to be true for Rho (Fig8C) and to be conserved for bovine GDI1 (SuppFig8). These results confirm that the GDI-QQ mutant is indeed extraction deficient: modestly so for inactive GTPases and completely so for active GTPases.

### RhoGDI extracts active RhoGTPase in vivo

We sought to directly test whether RhoGDI can extract active GTPases *in vivo* by employing GDI-QQ. First, we compared the effects of WT vs. QQ GDI overexpression on wounded oocytes expressing constitutively-active Cdc42 (G12V). (Q61L could not be used as it failed to elevate Cdc42 levels around wounds; see SuppFig11 and Supplemental Discussion). While WT GDI significantly reduced the amount of constitutively-active Cdc42 around wounds, GDI-QQ did not (Fig9A,B). Second, we compared the effects of WT vs. QQ GDI overexpression on wounded oocytes microinjected with Cy3-Cdc42 bound to GTPɣS. WT GDI significantly reduced the amount of Cy3-Cdc42(GTPɣS) around wounds while GDI-QQ did not (Fig9C,D). Collectively, these data suggest that GDI can extract active GTPase from the plasma membrane *in vivo*.

**Figure 9:**
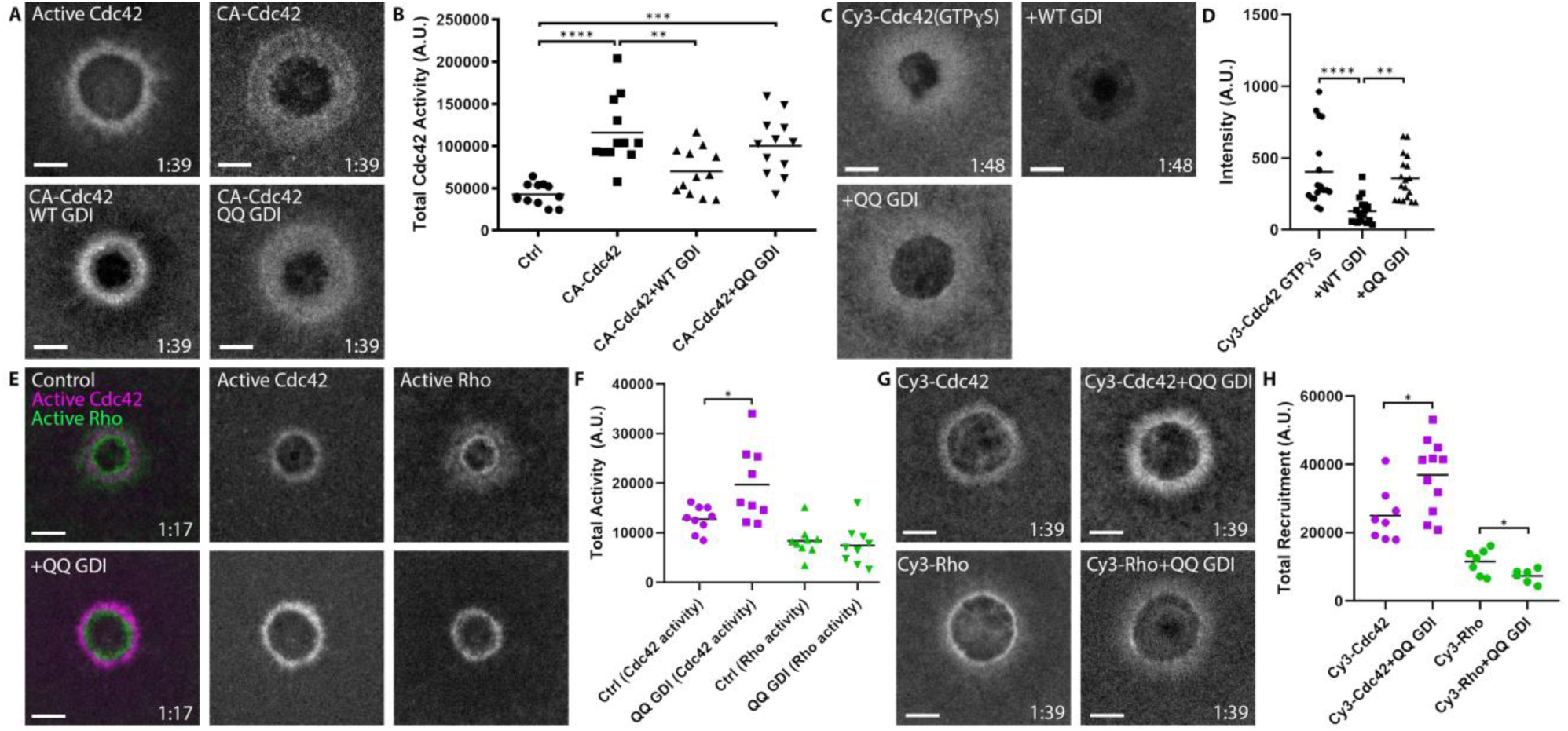
RhoGDI extracts active Cdc42 *in vivo*. A) Oocytes microinjected with wGBD alone or with constitutively-active Cdc42 (G12V), WT or QQ GDI; B) Quantification of total Cdc42 activity for (A), (n=12); C) oocytes microinjected with Cy3-Cdc42 bound to GTPɣS alone or with WT or QQ GDI; D) Quantification of intensity for (C), (n=18). Scale bar 10μm, time min:sec. One-way ANOVA with Tukey post-test statistical analysis; E) Oocytes microinjected with wGBD (magenta), rGBD (green) alone or with QQ GDI; F) Quantification of total Cdc42 (magenta) and Rho (green) activity from (E) (n=9); G) Cy3-Cdc42 or Cy3-Rho alone or with QQ GDI; H) Quantification of total recruitment of Cy3-Cdc42 (magenta) and Cy3-Rho (green) (n=6-11). Scale bar 10μm, time min:sec. Unpaired student’s t-test, 2-tailed distribution, unequal variance statistical analysis. *p<0.05, **p<0.01, ***p<0.001, ***p<0.0001.

The above results imply that the extraction of active RhoGTPase by GDI might contribute to its spatiotemporal patterning *in vivo*. To test this hypothesis, we expressed GDI-QQ and monitored the consequences on Rho and Cdc42 activity following wounding. Strikingly, Cdc42 activity around wounds was significantly elevated, in contrast to Rho which was unaffected (Fig9E,F). To assess whether the increase in activity was due to an increase of total Cdc42 around wounds as opposed to competition between GDI-QQ and the Cdc42 activity probe, we repeated the experiment with Cy3-Cdc42. Similar to the results obtained with the activity reporter, expression of GDI-QQ elevated Cy3-Cdc42 levels around wounds relative to controls (Fig9G,H). Cumulatively, these data suggest that GDI directly extracts active Cdc42 throughout the Cdc42 zone, and that extraction of active Cdc42 is necessary for its regulation around wounds.

## Discussion

Direct visualization of the RhoGTPases in living cells is essential for the understanding of their complex spatiotemporal dynamics. We have established two methods to fluorescently label vertebrate GTPases such that they are functional: internal tagging with a fluorescent protein or by sortase-mediated labeling with a fluorescent dye. This now provides us with widely applicable reagents to analyze GTPase function. These probes faithfully mimic the distribution of the endogenous GTPases based on their comparison to activity reporters in several processes: cell wound repair, cytokinesis, junctional integrity and epithelial wound repair. Further, the successful rescue of Rho function at wounds in the presence of C3 by a C3-insensitive mutant of IT-Rho indicates that these proteins are capable of replacing their endogenous counterparts. It will be important to assess the ability of IT- or Cy3-labeled GTPases substitute for their endogenous counterparts in other vertebrate cellular processes in the future, although we note that IT-Cdc42 has been shown to be functional in fission yeast (Bendezú et al., 2015). Nonetheless, the combination of the two labeling approaches is powerful as it permits side-by-side comparison of results obtained *in vivo* and *in vitro,* as demonstrated here.

Visualization of functional, labeled RhoGTPases in combination with activity reporters in living cells led to an unexpected observation: pools of inactive Cdc42 at the plasma membrane. To the best of our knowledge, this is the first time that inactive GTPases have been detected at membranes under conditions other than gross GTPase overexpression. Notably, the pool of inactive Cdc42 spatially coincides with a local Cdc42-GAP, Abr (Vaughan et al. 2011). The pool of inactive Cdc42 expands with overexpression of Abr, further demonstrating that locally-inactivated Cdc42 can remain associated with the plasma membrane. This finding has important mechanistic implications for the regulation of GTPase activity. It suggests that GTP hydrolysis and extraction of GTPases, while are likely indirectly linked, are not necessarily tightly coupled. This raises the possibility that GTPases might cycle through multiple rounds of activation and inactivation while remaining associated with the membrane.

Remarkably, in addition to the RhoGTPases themselves, GDI also localized to the plasma membrane in proximity to wounds. The localization of GDI in the same place where the GTPases are especially abundant implies that its accumulation reflects interaction with is GTPase clients. Consistent with this idea, GDI mutants deficient in GTPase binding fail to recruit to wounds (data not shown). It will be important to investigate the detailed mechanism of GDI localization and the control of its turnover at sites of high GTPase activity in the future.

The most significant result of this study is that RhoGDI extracts active GTPase, particularly active Cdc42, during cell wound repair. This finding is based on both *in vitro* assays showing that WT, but not GDI-QQ, extracts active GTPase from supported lipid bilayers and the *in vivo* demonstration that WT, but not GDI-QQ, extracts constitutively-active and GTPɣS-loaded Cdc42 from the plasma membrane. We thus conclude that GDI has the capacity to extract active GTPases. Moreover, this ability is harnessed to limit the level of Cdc42 activity during cell repair. While this finding may seem heretical, it has the virtue of explaining previous results in the oocyte wound repair system. That is, based on an indirect approach involving photoactivatable Rho and Cdc42 activity reporters, it was found that Cdc42 activity is lost throughout its zone, while Rho activity is preferentially lost at the trailing edge of its zone (Burkel, Benink, Vaughan, von Dassow, & Bement, 2012). The results presented here suggest that GDI is responsible for the removal of active Cdc42 throughout the Cdc42 zone while Rho is inactivated by a trailing edge GAP prior to extraction. This would also explain why a mild overexpression of GDI significantly reduced Cdc42 activity but had no effect on Rho activity: loss of active Cdc42 can be controlled at the level of GDI while Rho inactivation is controlled at the level of a GAP (SuppFig9).

The broader implications of RhoGDI’s ability to extract active GTPase are two-fold. First, the canonical GTPase cycle needs a new branch in which active GTPase can be directly extracted from the plasma membrane (Fig10). Further, because GDI binding strongly inhibits GTP hydrolysis and nucleotide exchange (Hart et al., 1992; Ueda, Kikuchi, Ohga, Yamamoto, & Takai, 2001), active GTPase may exist in its soluble form in complex with GDI. However, complementary evidence from biochemical and biological studies suggest that active GTPases are less stably bound to GDI compared to their inactive form (Hodgson et al., 2016; Slaughter, Das, Schwartz, Rubinstein, & Li, 2009; Tnimov et al., 2012). As such, the secondary extraction branch may actually represent a loop through which active GTPases are not only removed from cellular membranes, but rapidly returned to them (Fig10). Such a mechanism might enhance the spatial reach of GTPase activity within the plasma membrane or even mediate its spreading between different membrane compartments (Palamidessi et al., 2008).

**Figure 10:**
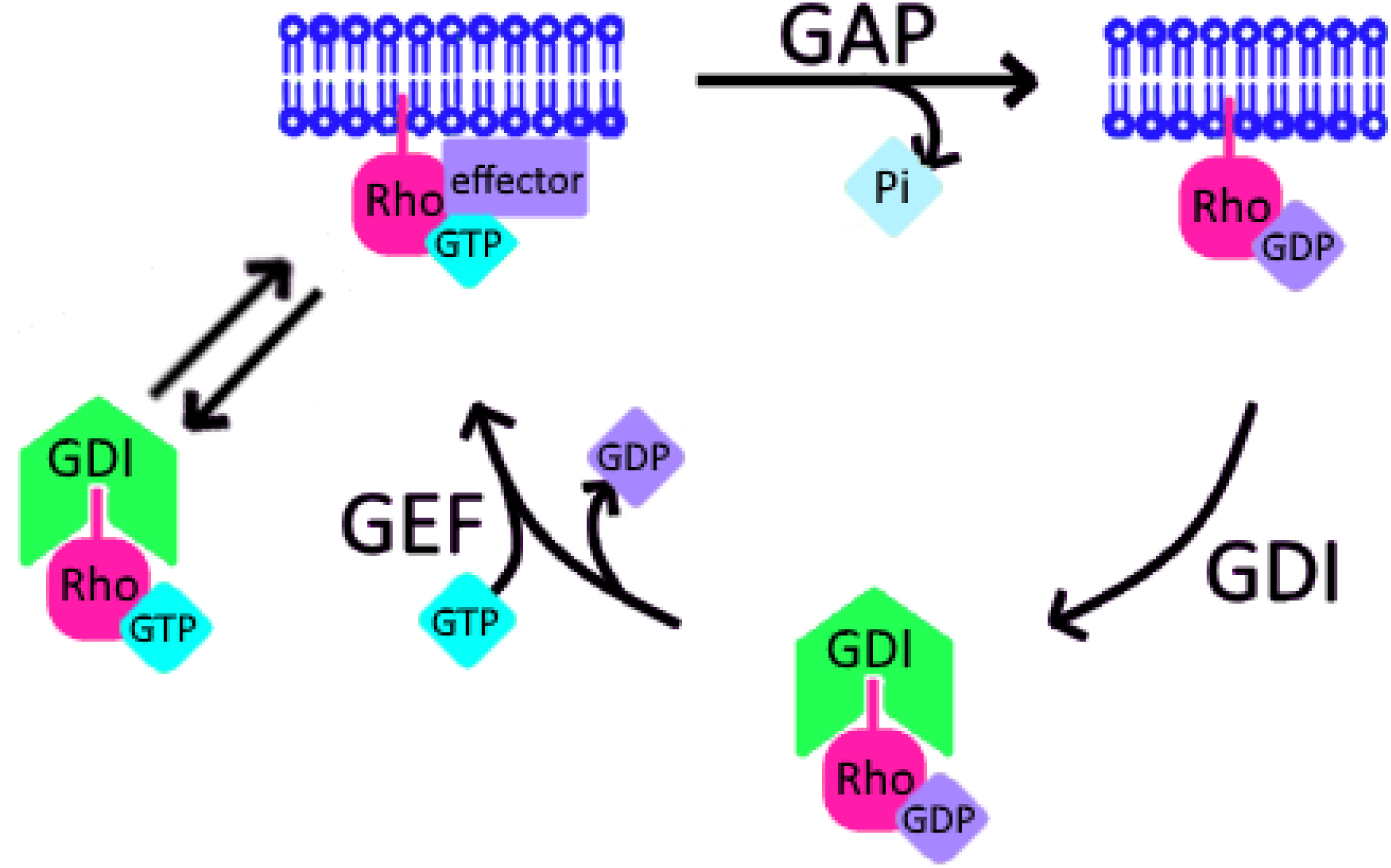
Schematic of proposed update to RhoGTPase cycle. We propose that in addition to the canonical GTPase cycle, GDI can extract active GTPase from the plasma membrane. Based on evidence that GTPase:GDI binding prevents GTP hydrolysis and nucleotide exchange (Hart et al., 1992; Ueda et al., 2001), active GTPase extracted by GDI would still be active upon its release back into the plasma membrane.

Second, RhoGDI’s ability to extract active GTPase forces us to reassess its role in GTPase regulation in different cellular processes. While the field primarily studies local GTPase regulation at the level of GEFs and GAPs, we should reconsider GDI’s role in regulation, as well as the regulation of GDI itself.

## Materials and Methods

### Plasmids

The active RhoGTPase probes, mRFP-wGBD, eGFP-wGBD, eGFP-2xrGBD, BFP-2xrGBD and mRFP-2xrGBD in pCS2+ were generated as previously described (Sokac et al. 2003; Benink and Bement 2005; Davenport et al. 2016). mCh-Rho, mCh-Rac, mCh-Cdc42 (Benink 2005), untagged Rho, Rac, and Cdc42 (wild-type (WT) and constitutively-active (G12V)) in pCS2+ were made as previously described (Benink and Bement 2005). For expression and purification in *E. coli*, codon-optimized Cdc42 and RhoA (Eurofins Genomics Germany GmbH, Ebersberg, Deutschland) were subcloned into a pETMz2 vector via Gibson assembly cloning (Gibson et al., 2009), with primers GTPase(GeneStrand)fwd and -rev and pETfwd and -rev (all primer sequences in SuppTable1). A pentaglycine for sortase-mediated labeling was added onto the 5’ of the GTPases. The constitutively-active Cdc42 G12V and RhoA G14V mutants were generated by Quickchange mutagenesis with primers Cdc42(G12V)fwd and -rev and RhoA(G14V)fwd and -rev, respectively.

*X. laevis* IT-Cdc42 in pCS2+ was generated according to Bendezú et al. (2015): a linker - SGGSACSGPPG- was cloned into Cdc42 after Q134. The linker encodes for *BamH1* and *Asc1* restriction sites for digestion and insertion of GFP into the linker region. The 5’ end of Cdc42 was amplified with primers Cdc42(1) and Cdc42(2); the 3’ end was amplified separately with primers Cdc42(3) and Cdc42(4). The two products were joined by PCR stitching with primers Cdc42(1) and Cdc42(4). The single product was digested with *EcoR1* and *Xho1* and ligated into pCS2+. The resulting construct was mutated by Quickchange with primers pCS2+-Cdc42(1) and pCS2+-Cdc42(2) to remove the *BamH1* restriction site upstream of the insertion in the multiple cloning site. eGFP was amplified from eGFP-wGBD with primers eGFP(1) and eGFP(2). Both the Quickchanged construct and eGFP were digested with *BamH1* and *Asc1*, and eGFP was ligated into the linker region internal to the Cdc42 coding sequence. *X. laevis* Rho and Rac were similarly tagged internally after residues Q136 and L134, respectively.

*X. laevis* RhoGDI Clone ID:7010361 (GE-Healthcare Dharmacon, Lafayette, CO) was subcloned into pCS2+ with *Cla1* and *Xho1*, and into N’3xGFP and N’Halo (Promega, Madison, WI) pCS2+ with *BspE1* and *Xho1*. A FLAG-tag was added by PCR onto the 5’ of RhoGDI, and the product was subcloned into pFast-Bac1 with *Cla1* and *Not1*. The following mutations were made by Quickchange mutagenesis to untagged and N’3xGFP RhoGDI in pCS2+: E158/9A, E158/9Q, D40A, D40N, D180A and D180N (Dransart et al., 2005). Mutant RhoGDI 8(A) had the first 8 charged amino acids to mutated to alanine (D3/5, E11-13, E15-17A) by sequential PCR (Ueyama et al., 2013). The 3’ end of RhoGDI was amplified with primers 8(A)F1 and R1. The product was amplified and added to at its 5’ end with primers 8(A)F2 and R1, and for a third time with 8(A)F3 and R1. The final PCR product was subcloned into pCS2+ by Infusion PCR (Takara Bio, Kusatsu, Japan). For subcloning into N’3xGFP-pCS2+, the third PCR from above was repeated with 8(A)F4 and R2. Mutant RhoGDI helix replacement (HR) had alpha helix D39-Q48 replaced with a glycine linker GGGGSGGGGS. This was done by sequential PCR as described above with four rounds of PCR: HR1 and R1, HR2 and R1, HR3 and R1, then either HR4 and R1 for subcloning into pCS2+ or HR5 and R2 for subcloning into N’3xGFP-pCS2+. RhoGDI mutant Δ51-199 was generated by adding a stop codon after L50 by Quickchange mutagenesis. To make RhoGDI Δ1-19, the 3’ end of RhoGDI was amplified with primers (-)20 F1 and R for subcloning by infusion into pCS2+, and primers (-)20 F2 and R for N’3xGFP-pCS2+. Primers (-55) F1, F2 and R were used to generate RhoGDI Δ1-54 as described above (Hoffman et al., 2000; Ueyama et al., 2013). For expression and purification in *E. coli*, *X. laevis* RhoGDI WT and E158/9Q were subcloned into a pGEX-6P-2 vector via Gibson assembly cloning (Gibson et al., 2009) using XlRhoGDIfwd and -rev. A cysteine for labeling was added by PCR onto the 5’ of RhoGDI. Bovine RhoGDI1 in pGEX-6P was a gift of Dr. Tomotaka Komori. E163/4Q mutation was made by Quickchange mutagenesis with the E163/4Qfwd and - rev primers. Mutant bovine RhoGDI Δ1-22 and Δ1-59 were subcloned with BamHI and NotI into a pGEX-6P-2 with Δ1-22fwd and -rev and Δ1-59fwd and -rev, respectively. Mutant bovine RhoGDI HR was generated via Gibson assembly cloning (Gibson et al., 2009), with primers HRfwd and -rev and pGEXHRfwd and -rev, for amplification of RhoGDI and the pFASTBacH10 vector, respectively. RabGGTase 2 beta in a pGATEV vector was kindly provided by Dr. Konstantin Gavriljuk.

### Expression and purification of recombinant protein from E. coli for sortase-labeling

Rosetta(DE3) chemically competent *E. coli* cells were transformed with WT or mutant RhoGDIs, induced with 250µM IPTG and incubated at 18°C ON. Bacteria cells were harvested, centrifuged at 4000xg for 20min, and pellets flash frozen in liquid nitrogen and stored at -80°C. Frozen pellets were resuspended in a 3x volume of lysis buffer (50mM KPi pH 8, 400mM KCl, 1mM EDTA, 5mM βME, 1mM PMSF, 1mM benzamidine) and lysed with a high pressure homogenizer at 4°C. Lysate was clarified by centrifugation at 100,000xg for 1hr and applied to a glutathione sepharose 4 fast flow column bed (GE-Healthcare, Chicago, IL) equilibrated with wash buffer (50mM KPi pH 8, 400mM KCl, 1mM EDTA, 5mM BME, 1mM benzamidine). The column was washed with wash buffer, and protein was eluted with elution buffer (50mM KPi pH 8, 400mM KCl, 1mM EDTA, 5mM BME, 1mM benzamidine, 10mM reduced L-glutathione). Peak fractions were pooled, protein concentration was estimated with Bradford assay (Bio-Rad Laboratories, Inc., Hercules, CA), and PreScission protease was added at 1:30. After ON incubation on ice, the sample was concentrated using 5,000 MWCO Vivaspin15R centrifugal concentrators (Sartorius AG, Göttingen, Germany), buffer exchanged in wash buffer on a HiPrep 26/10 desalting column (GE-Healthcare) and recirculated on the same glutathione sepharose 4 fast flow column bed re-equilibrated in wash buffer. Flow through was collected, concentrated, spun down and gel filtered on a HiLoad Superdex 75 pg column (GE-Healthcare) in storage buffer (20mM HEPES pH 7.5, 150mM KCl, 0.5mM TCEP, 20% Glycerol). Peak fractions were pooled, concentrated, flash frozen in liquid nitrogen and stored at -80°C. Protein purification and purity were determined by Coomassie stain of 12% SDS-PAGE, protein concentration measuring absorbance at 280nm. WT and mutant GTPases were expressed and purified similarly to GDIs with the following differences: L21(DE3) chemically competent *E. coli* cells were used and proteins were expressed with 1 mM IPTG at 37°C for 4hrs. The affinity step was performed on HiTrap Chelating HP columns loaded with cobalt and equilibrated in 50mM HEPES pH 7.5, 50mM NaCl, 5mM MgCl2, 0.5mM BME, 100µM ATP and 100µM GDP/GTP. RabGTTase Beta was expressed and purified as described before (Gavriljuk, Itzen, Goody, Gerwert, & Kötting, 2013). After gel filtration, nucleotide bound to Cdc42 G12V was exchange to GTPγS, incubating the protein with 10-fold excess EDTA and GTPγS for 30 minutes on ice. The new nucleotide state was stabilized by adding 20-fold excess MgCl_2_. Before freezing, protein buffer was exchanged with a NAP-5 column (GE-Healthcare) in storage buffer (50mM HEPES pH 7.5, 50mM NaCl, 2mM MgCl2, 2mM DTT, 20% Glycerol).

### Cy3-labeling and in vitro prenylation of RhoGTPases

RhoGTPases were labeled at the N-t with Cy3 using a sortase-mediated reaction and *in vitro* prenylated as previously described (Gavriljuk et al., 2013; Popp, Antos, Grotenbreg, Spooner, & Ploegh, 2007). In brief, RhoGTPases were incubated with sortase and Cy3 N-t labeled LPETGG peptide at 3:1:15 ratio in labeling buffer (Tris pH 8.0, 150mM KCl, 6µM CaCl2, 0.5mM TCEP) and incubated ON at 16°C. The entire reaction was mixed with geranylgeranyltransferase type 1 and geranylgeranyl diphosphate at 10:1:30 ratio in prenylation buffer (50mM HEPES pH 7.5, 50mM NaCl, 2mM MgCl_2_, 2mM DTT, 30µM GDP/GTP, 2%

CHAPS), and incubated ON on a rotating mixer at 4°C. The sample was spun in a TLA-100 rotor (Beckman Coulter, Brea, CA) at 80,000 rpm for 30 minutes at 4°C and gel filtered on a HiLoad Superdex 75pg column (GE-Healthcare) equilibrated with prenylation buffer with 0.5% CHAPS. Peak fractions were pooled, concentrated using 5,000 MWCO Vivaspin4 centrifugal concentrators (Sartorius AG) and buffer exchanged in prenylation buffer without CHAPS on a NAP-5 column (GE-Healthcare). Residual detergent was removed by Pierce™ Detergent Removal Spin Column (Thermo Fisher). After sortase-mediated labeling with Cy3, unprenylated proteins were directly spun down and gel filtered in absence of CHAPS.

### Oocyte collection and preparation

Ovarian tissue was harvested from adult *X. laevis* via surgical procedures approved by the University of Wisconsin-Madison Institutional Animal Care and Use Committee. Oocytes were stored in 1x modified Barth’s solution (88mM NaCl, 1mM KCl, 2.4mM NaHCO_3_, 0.82mM MgSO_4_, 0.33mM NaNO_3_, 0.41mM CaCl_2_, 10mM HEPES, pH 7.4) with 100µg/mL gentamicin sulfate, 6µg/mL tetracycline and 25µg/mL ampicillin at 16°C. Prior to manual defolliculation with forceps, oocytes were treated with 8mg/mL type I collagenase (Life Technologies, Grand Island, NY) in 1x modified Barth’s solution for 1hr at 16°C on an orbital shaker.

### mRNA preparation

mRNA was generated *in vitro* using the mMessage mMachine SP6 transcription kit (Thermo Fisher, Carlsbad, CA) and purified using the RNeasy Mini Kit (Qiagen, Hilden, Germany). Transcript size was verified on a 1% agarose/formaldehyde denaturing gel relative to the Millennium Marker (Life Technologies) RNA molecular weight standard.

### Oocyte microinjection

Oocytes were microinjected with a 40nL injection volume using a p-100 microinjector (Harvard Apparatus, Holliston, MA). mRNA encoding probes for active Rho (2xrGBD) and active Cdc42 (wGBD) were injected at a final needle concentration of 30µg/mL and 100µg/mL, respectively. IT-Rho, Rac and Cdc42 were each injected at a final needle concentration of 125µg/mL, with 63µg/mL of WT RhoGDI to stabilize the exogenous GTPase and maintain stoichiometric ratio of GTPase:GDI (Boulter et al., 2010). mRNA encoding Abr was injected at a final needle concentration of 25-500µg/mL. 3xGFP-WT GDI and mutants at 333µg/mL, untagged RhoGDI at 300µg/mL, Halo-WT GDI and mutants at 200µg/mL, Cdc42 G12V at 28µg/mL, and untagged GDI E158/9Q at 1.5mg/mL. For purified protein, Cy3-Rho and Cy3-Cdc42, *in vitro* prenylated and complexed with RhoGDI, were injected at a final needle concentration of 4.56µM. C3 exotransferase was injected at a final needle concentration of 1.1µg/mL in 1mM DTT and WT GDI at 3.5-114µM for the standard curve. For wounding experiments, all mRNA was injected 20-24hrs before imaging, and purified protein was injected at least 2hrs before imaging, except for C3 which was injected 30min prior to imaging. For imaging cortical granule exocytosis, oocytes were injected 16hrs before imaging and matured ON in progesterone. Two-cell embryos were microinjected with a 5nL injection volume at a final needle concentration of 167µg/mL for IT-Rho and IT-Cdc42 mRNA, and 18.24µM Cy3-Rho and Cy3-Cdc42.

### Purification of recombinant protein from E. coli for antibody purification

BL21 pLysS cells (Thermo Fisher) were transformed with GST-RhoGDI in pGEX6p.1. A positive clone was used to inoculate 12mL of lysogeny broth (LB) supplemented with 25µg/mL ampicillin and cultured ON. The 12mL culture was added to 1L of LB with ampicillin and shaken at 37°C until OD600∼0.6. The culture was induced by adding a final concentration of 0.1mM IPTG and shaken at 37°C for 2hrs. BL21 pLysS cells were pelleted at 5300rpm for 10min at 4°C, and the pellet resuspended in Buffer A (50mM Tris-HCL, pH 7.6; 50mM NaCl with 1mM DTT in PBS). Pellets were stored at -80°C. Pellets were thawed at room temperature to promote cell lysis. Triton X-100 was added to a final concentration of 0.6%, PMSF at 500uM, lysoszyme at 1mM in 10mM Tris pH 8.0, 400µM Peflabloc, 1µg/mL aprotinin, 1µg/mL leupeptin. Solubilate was incubated at RT for 30min, DNAse1 was added to a final concentration of 10ug/mL, incubated again for at RT for 30min, and centrifuged at 16,000xg for 10min at 4°C. The supernatant was collected and exposed to a column containing glutathione-sepharaose 4B (MilliporeSigma). The column was washed 5x with Buffer A and the protein eluted with 20mM Tris, pH 8.0, 20mM glutathione, 400µM Peflabloc, 1.25µg/mL aprotinin, 14.25µg/mL leupeptin, 0.25mM E-64, 0.5mM PMSF. Protein concentration was determined by Coomassie stain of a 12% SDS-PAGE alongside a BSA standard curve.

### Antibody generation and purification

FLAG-RhoGDI purified from Sf9 cells was used as an antigen for antibody production in rabbits (Covance, Princeton, NJ). The serum was heat-inactivated at 56°C for 30min, diluted 1:1 in 20mM Tris, pH 7.5, and filtered through a 0.22µm syringe. The diluted, filtered serum was loaded onto a column containing GST-GDI coupled to Affi-Gel 15, to minimize antibody cross-reactivity to the FLAG-tag on the antigen. The column was washed 20x with 20mM Tris, pH 7.5 and 20x with 20mM Tris, pH 7.5, 500mM NaCl. Antibody was first eluted with 100mM glycine, pH 2.5 into 1M Tris, pH 8.8 for neutralization. The column was washed 20x with 20mM Tris, pH 8.8. Antibody remaining on the column was eluted with 100mM Triethylamine, pH 11.5 into concentrated HCl and 1M Tris, pH 7.5 for neutralization. The concentration of each fraction was determined by A_280._ The peak antibody fractions were pooled, dialyzed against PBS (2x2L) ON, and concentrated using a 100K MW Amicon Ultra-15 Centrifugal filter (MilliporeSigma). Antibody specificity was determined by western blotting of purified protein, *X. laevis* oocyte whole cell lysate (WCL), WCL of oocytes overexpressing GDI and WCL of oocytes expressing 3xGFP-GDI.

### Expression and purification of recombinant protein from insect cells

DH10Bac-competent *E. coli* (Thermo Fisher) were transformed with FLAG-WT RhoGDI or E158/9Q in pFast-Bac1 and positive clones were identified by blue/white screening. Bacmid was purified and transfected into Sf9 cells using Cellfectin II reagent (Thermo Fisher). High-expressing clones were identified and baculovirus was generated for two additional passages. Sf9 cells, 22x10^6^ per 15cm plate, were infected with high-titer baculovirus and incubated 27°C for 72hrs. Sf9 cells were harvested, centrifuged at 500xg for 5min, and pellets were stored at -80°C.

Frozen pellets were resuspended in a 5x volume of solubilization buffer (1xPBS pH 7.5, 1% Triton X-100, 0.5µg/mL leupeptin, 0.5µg/mL aprotinin, 0.5µg/mL Pepstatin A, 40µg/mL PMSF, 100µg/mL benzamidine, 0.5µg/mL E64) and incubated at 4°C with end-over-mixing for 1hr. Lysate was clarified by centrifugation at 21,000xg for 15min and applied to an anti-FLAG M2 agarose column bed (MilliporeSigma, Burlington, MA). The column was washed 3x with wash buffer (1xPBS, 0.5µg/mL leupeptin, 0.5µg/mL aprotinin, 0.5µg/mL Pepstatin A, 40µg/mL PMSF, 100µg/mL benzamidine, 0.5µg/mL E64). A buffer exchange was performed with 1:1 wash buffer:HEPES (25mM HEPES pH 7.5, 100mM KCl), and the column washed 2x with HEPES. Protein was eluted with 1M Arginine pH 4.4 into an equal volume of collection buffer (50mM HEPES pH 7.5, 200mM KCl). Fractions were analyzed by coomassie stain of a 12% SDS-PAGE. Peak fractions were pooled and concentrated using a 10MW Amicon Ultra-15 Centrifugal filter (MilliporeSigma). A buffer exchange was performed during concentrating with HEPES such that the final Arginine concentration was less than 1mM. Protein purification and purity was determined by comparison to a BSA standard curve by Coomassie stain of a 12% SDS-PAGE.

### Fixing and staining of wounded oocytes

Oocytes were wounded, allowed to heal for 2-3min and fixed for 2hrs in 10mM EGTA, 100mM KCl, 3mM MgCl_2_, 10mM HEPES, 150mM sucrose (pH 7.6), 4% PFA, 0.1% glutaraldehyde, 0.1% Triton X-100. Fixed oocytes were washed 5x in TBSN/BSA (5mg/mL BSA in 1xTBS containing 0.1% NP-40). Oocytes were bisected and blocked in TBSN/BSA for 4hrs at 4°C. Oocytes were stained with rabbit α-RhoGDI at 1:1000 in TBSN/BSA for 12hrs, washed 5x in TBSN/BSA over 12hrs, stained with chicken α-rabbit Alexa Fluor 647 (Invitrogen, Carlesbad, CA) at 1:10,000 for 12hrs in BSN/BSA at 4°C, and washed 5x in TBSN/BSA over 12hrs.

### Image acquisition, wounding and data analysis

Laser scanning confocal microscopy was performed using a Nikon Eclipse Ti inverted microscope with a Prairie Point Scanner confocal system (Bruker, Middleton, WI). The microscope was fitted with a 440-nm dye laser pumped by a MicroPoint 337-nm nitrogen laser (Andor, South Windsor, CT) for wounding. Brightest-point projections, measurements of fluorescence intensities, area and distances were made in FIJI (Schindelin et al., 2012). Bio-Formats Importer and De-Flicker plugins were used. Ring intensity corrected for background was calculated by quantifying the mean intensity of the ring and subtracting the mean intensity of the background. Total activity was calculated by multiplying the mean intensity of the zone (corrected for background) by the area of the zone, normalized for wound width. GraphPad Prism was used to plot quantifications and perform statistical analyses. An unpaired student’s T-test with a 2-tailed distribution and unequal variance was used to compare two conditions, one-way ANOVA with a Tukey post hoc analysis was used to analyze more than two conditions. *p<0.05, **p<0.01, ***p<0.001, ****p<0.0001.

### Supported lipid bilayer assay

Small unilamellar vesicles and supported lipid bilayers were prepared with 100% 1,2-dioleoyl-sn-glycero-3-phosphocholine (18:1 DOPC; Avanti Polar Lipids, Inc., Alabaster, AL) as described before (Hansen et al., 2019). 250µL Cy3 labeled GTPases were incubated on SLBs at 200nM final concentration until equilibrium was reached. A 0.5mm silicone tubing was attached drop-to-drop to the chamber with the equilibrated sample via a male luer connector (ibidi GmbH, Gräfelfing, Germany). After acquisition of few frames in absence of flow, the chamber was flushed at 10µL/sec with imaging buffer (20mM HEPES pH 7.0, 150mM KCl, 1.5mM MgCl_2_, 0.5mM EGTA, 100µM GDP/GTP) alone or in presence of a GTPase solubilizer until baseline was reached. Oxygen scavenger system (1.25mg/mL glucose oxidase, 0.2 mg/mL catalase, 400 mg/mL glucose) was added fresh to each sample and buffer before imaging. TIRF was performed on a Nikon Eclipse Ti inverted microscope with a VisiScope TIRF-FRAP Cell Explorer system (Visitron Systems GmbH, Puchheim, Germany) using a 60× Apo TIRF oil-immersion objective (1.49 N.A.). Cy3-labeled proteins were excited with a 561-nm laser line, excitation light was passed through a ET-561nm Laser Bandpass Set (Chroma Technology Corporation, Bellows Falls, VT) before illuminating the sample. Fluorescence emission was detected on a Evolve 512 Delta EMCCD camera (Teledyne Photometrics, Tucson, AZ). Measurements of fluorescence intensities were made in FIJI (Schindelin et al., 2012). Plot Z-axis profile tool and Bio-Formats Importer plugins were used. Fluorescence intensity was corrected for background. To display multiple curves on the same graph, data from different experiments were aligned using the overshoot signal occurring after the flow was started and normalized dividing by the maximum intensity. Data from wash off experiments were fitted with a one component exponential decay function (*y* = *y*_0_ + *A e*^−λ *x*^; *y*_0_ = y offset, *A*=amplitude, λ=exponential decay constant), choosing a fitting range that did not include the initial overshoot. This was possible because a monoexponential function can be fitted to a range of the data set without affecting the λ value obtained. K_off_ titration curves were fitted with a hyperbolic function 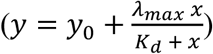. Origin Pro (OriginLab Corporation, Northampton, MA) was used to analyze data, plot quantifications and perform statistical analyses. An unpaired student’s T-test with a 2-tailed distribution and equal variance was used to compare two conditions. **p<0.01, ***p<0.001, ****p<0.0001.

## Acknowledgements

This work was supported by National Institutes of Health Grant GM52932 to W.B., a Dr. Stanley and Dr. Eva Lurie Weinreb Fellowship to A.G. and HSFP CDA00070/2017-2 to P.B. I.V is supported by the MaxSynBio Consortium, which is jointly funded by the Federal Ministry of Education and Research of Germany and the Max Planck Society. We acknowledge National Institutes Health Grant R44 MH065724 to LOCI at UW-Madison. We are also grateful to both our labs for their continued input.

---

**Supplemental Figure 1:**
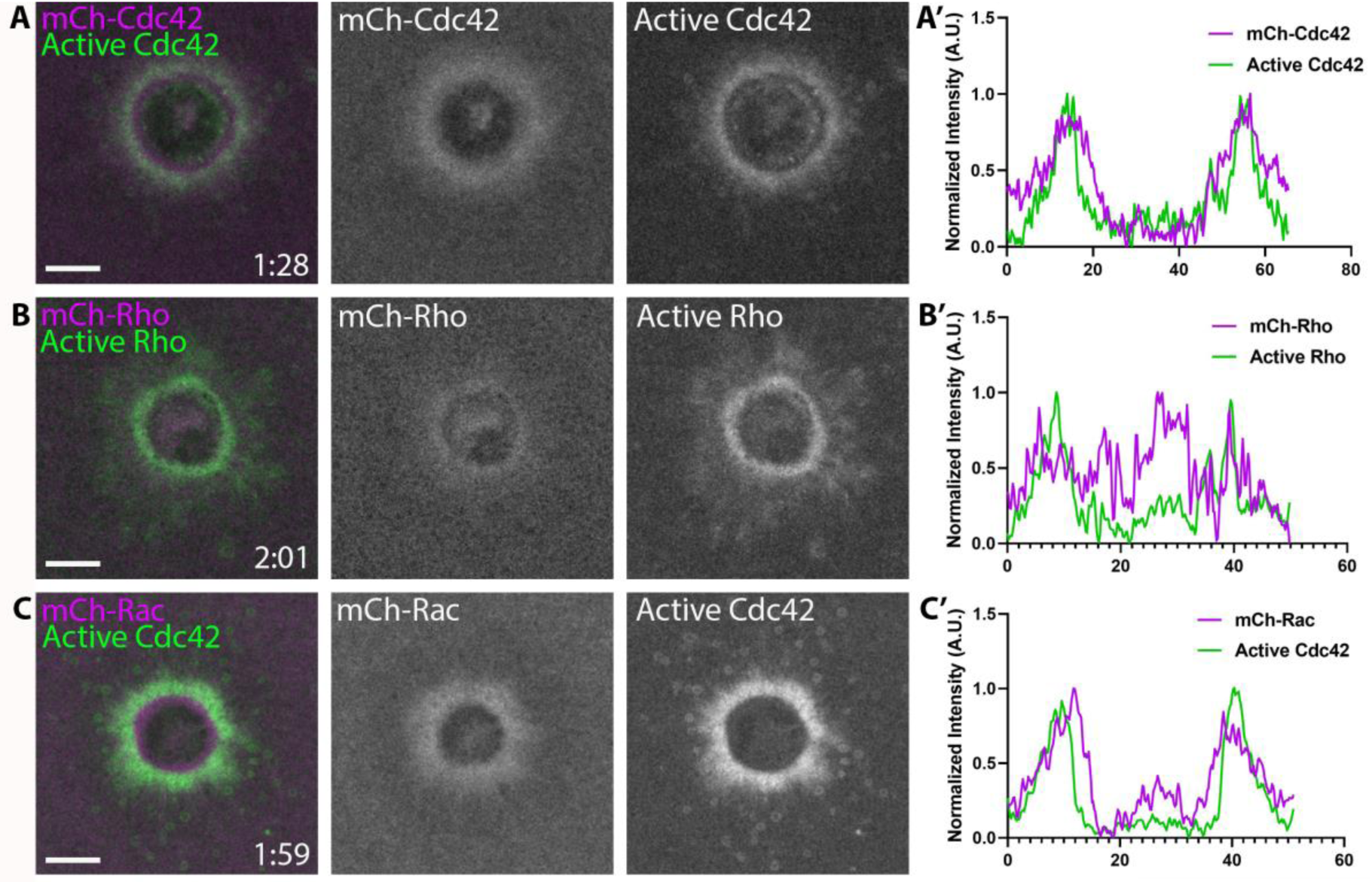
Amino-terminally tagged RhoGTPases do not localize properly to wounds. Oocytes injected with A) mCh-Cdc42 (magenta) and wGBD (green), B) mCh-Rho (magenta) and rGBD (green) or C) mCh-Rac (magenta) and wGBD (green) with A’-C’) Corresponding line scans. Scale bar 10μm, time min:sec.

**Supplemental Figure 2:**
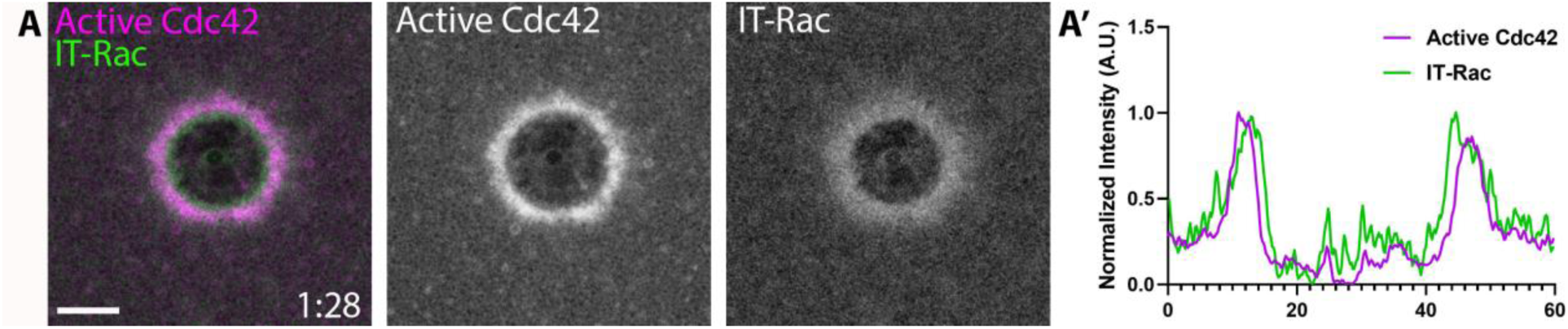
Internally-tagged Rac localizes to wounds. A) Oocyte injected with wGBD (magenta) and IT-Rac (green); A’) Corresponding line scan. Scale bar 10μm, time min:sec.

**Supplemental Figure 3:**
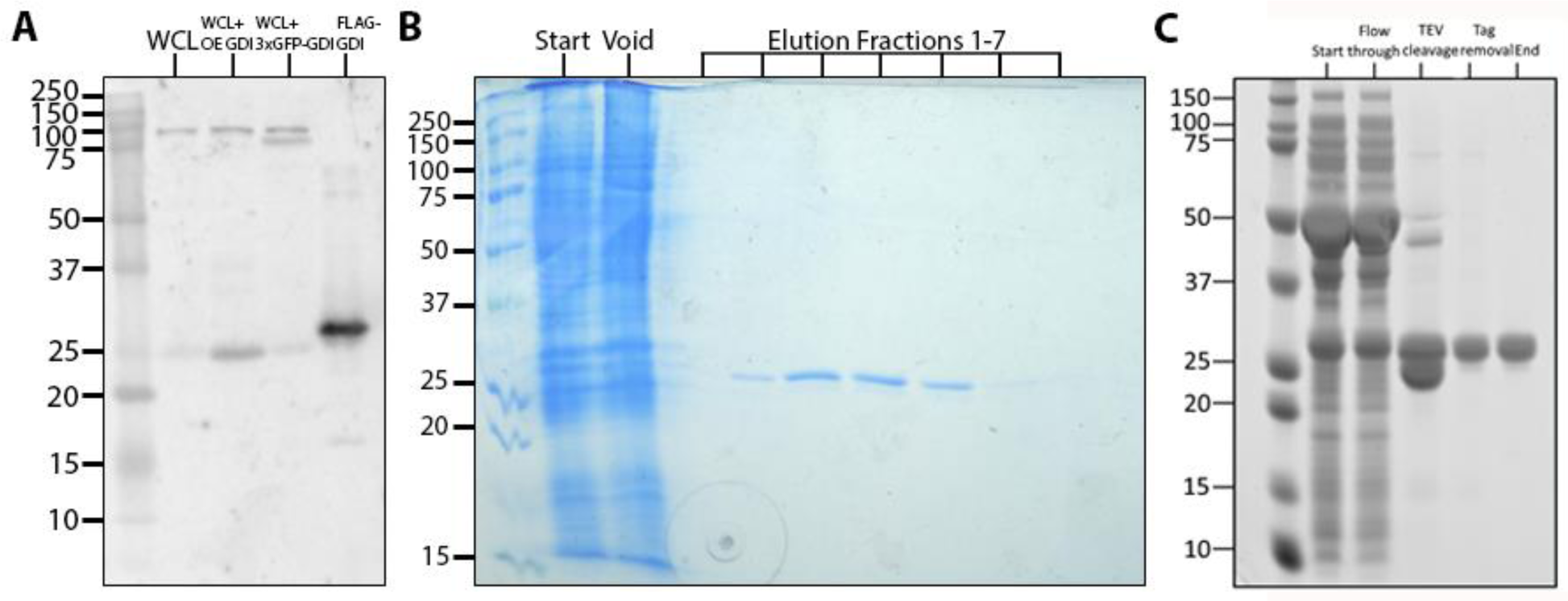
*X. laevis* RhoGDI antibody specificity and purified RhoGDI protein. A) Western blot stained with αGDI antibody to determine specificity; lane 1: X. laevis oocyte whole cell lysate (WCL), lane 2: WCL of oocytes overexpressing GDI, lane 3: WCL of oocytes expressing 3xGFP-RhoGDI, lane 4: purified FLAG-RhoGDI; B) Coomassie stain of 12% SDS-PAGE to assess purity of FLAG-RhoGDI; lane 1: start, lane 2: void, lanes 3-10: elution fractions. C) Coomassie stain of 12% SDS-PAGE to assess purity of RhoGDI purified from bacteria; lane 1: start, lane 2: flow through, lane 3: TEV cleavage, lane 4: tag removal, lane 5: end product after gel filtration.

**Supplemental Figure 4:**
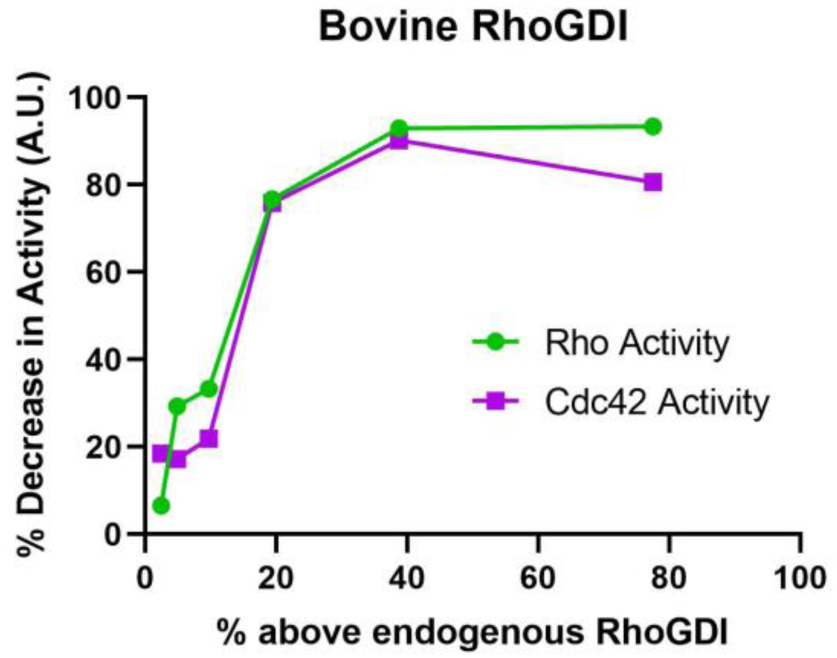
Bovine RhoGDI decreases Rho and Cdc42 activity in a dose-dependent manner *in vivo*. Standard curve of decrease in Rho and Cdc42 activity with increasing concentrations of bovine RhoGDI (n=2-30 for each concentration).

**Supplemental Figure 5:**
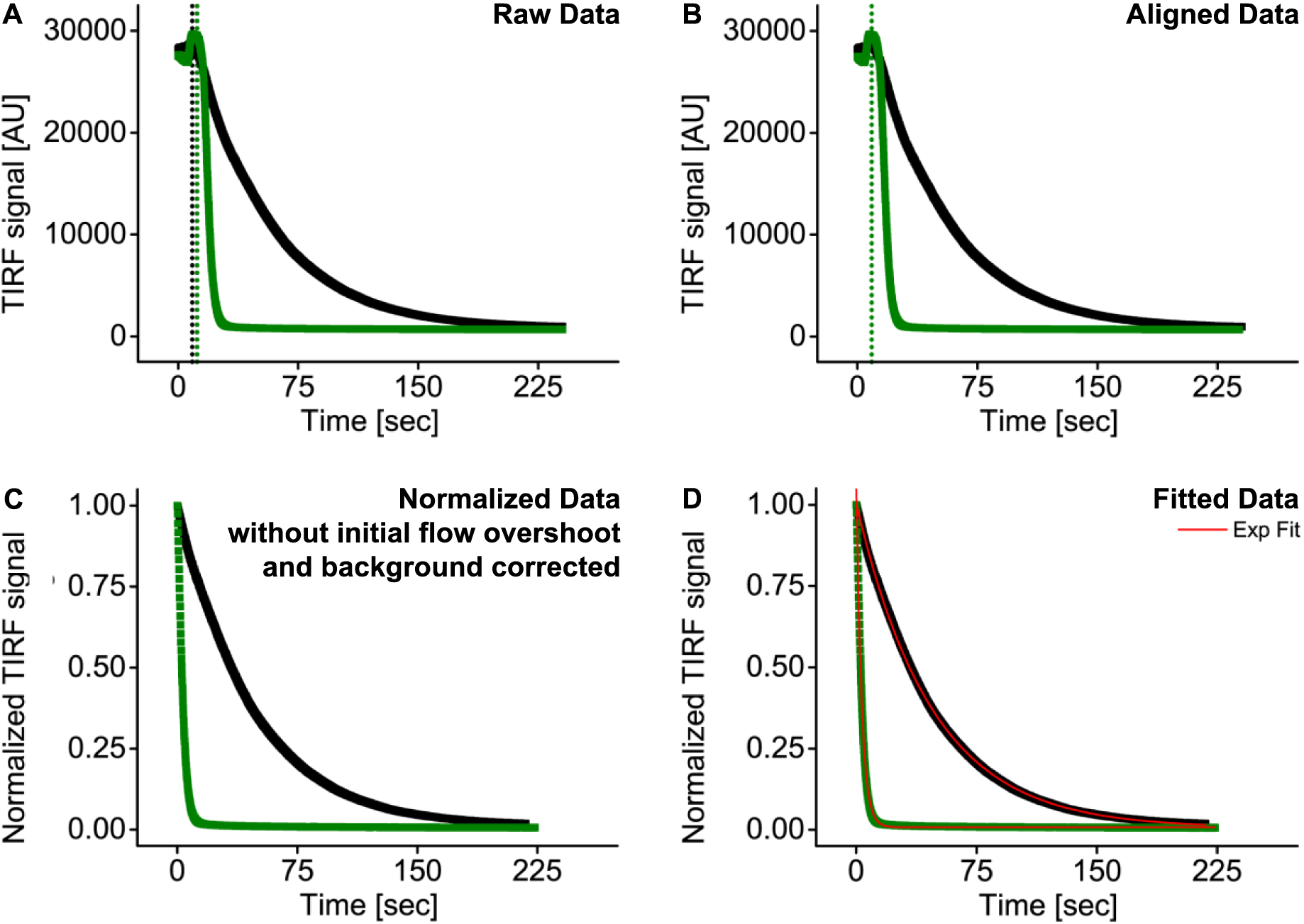
*In vitro* data analysis. A) Raw data. Wash off experiments were started after the RhoGTPase signal at the membrane was stable. After starting the syringe pump, a signal overshoot occurred; B) The overshoot signal was used as a reference point to align different experiments and cut off for further analysis; C) Data were background corrected and normalized to display multiple curve on the same graph; D) Data were fitted with a monoexponential decay curve to obtain dissociation constants (K_off_) of the GTPases from the membrane.

**Supplemental Figure 6:**
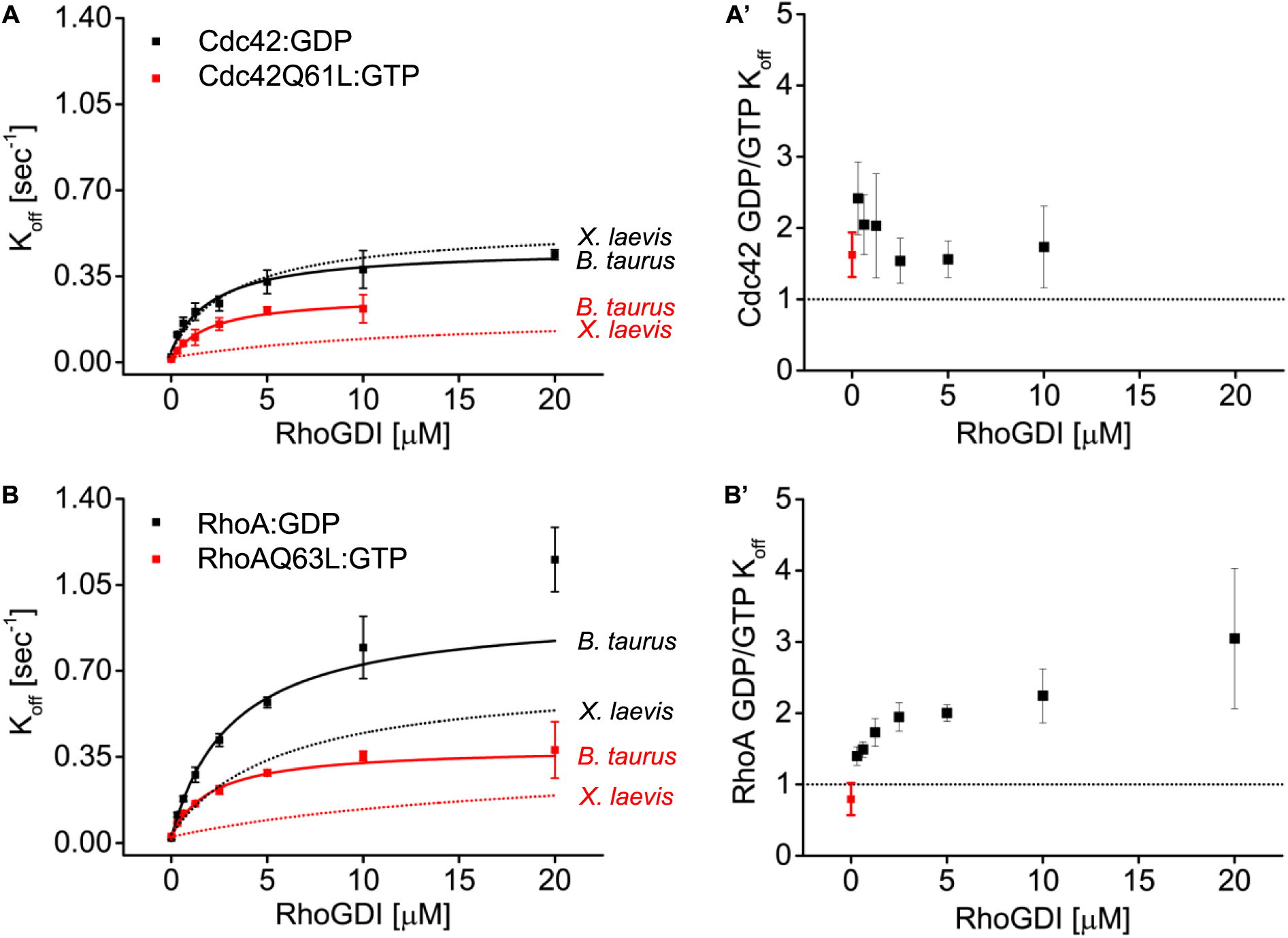
Comparison of bovine and *Xenopus* RhoGDI in their ability to actively extract both inactive and active RhoGTPases from synthetic membranes. A) K_off_ values obtained for inactive and constitutively active Cdc42 at different bovine RhoGDI concentrations. Extraction rates were fitted with a hyperbolic function; A’) Ratio of K_off_ obtained for inactive and constitutively active Cdc42 at the same bovine RhoGDI concentration; B-B’) same as in A-A’ for inactive (RhoA:GDP) and constitutively active (RhoAQ63L:GTP) RhoA.

**Supplemental Figure 7:**
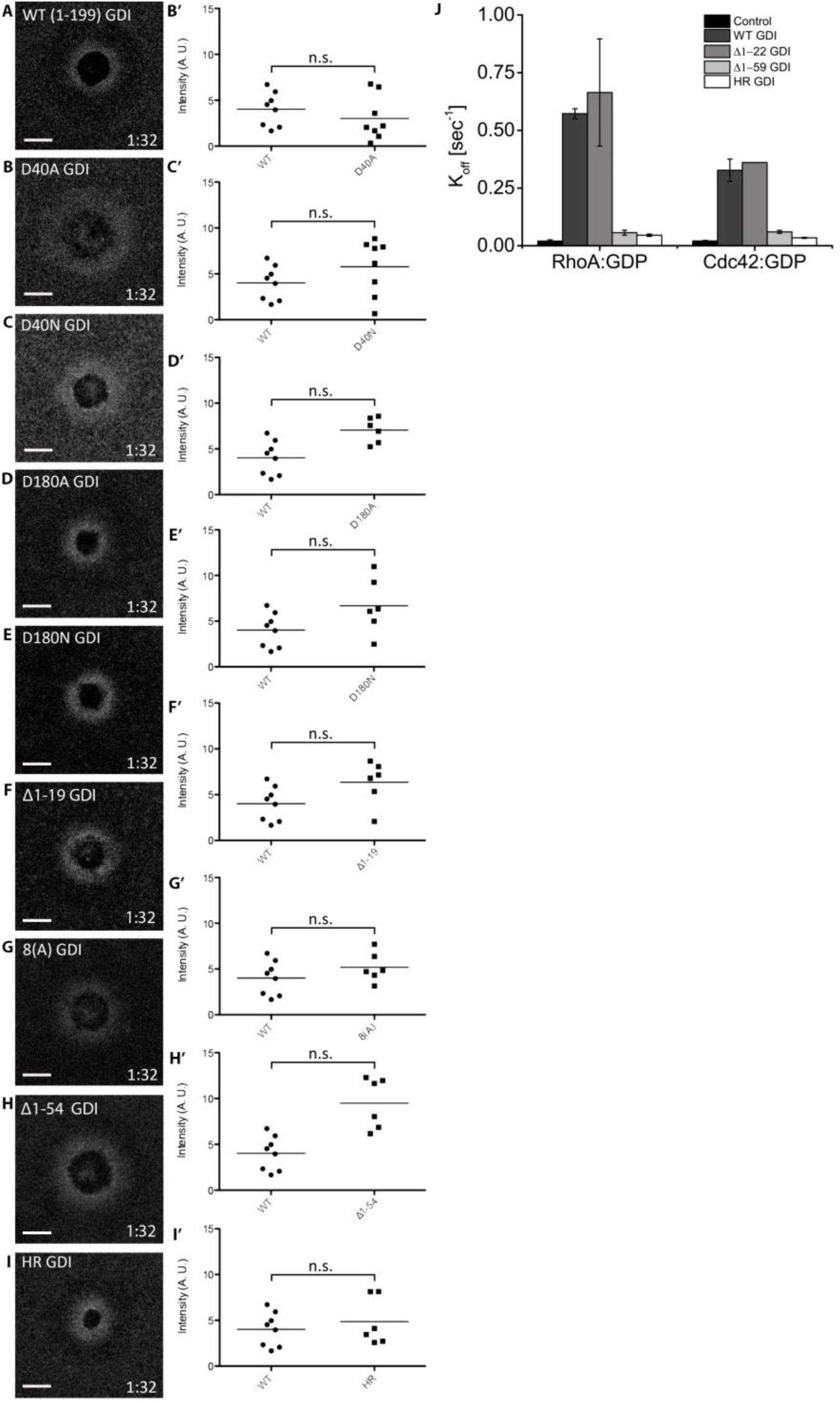
Analysis of previously-described extraction-deficient mutants. A-I) Oocytes microinjected with WT or mutant RhoGDI and quantification of localization relative to WT RhoGDI; Unpaired student’s t-test, 2-tailed distribution, unequal variance statistical analysis; J) Average K_off_ values obtained for inactive RhoA and Cdc42 in absence (control) and presence of 5 μM of either bovine RhoGDI WT or bovine RhoGDI mutants.

**Supplemental Figure 8:**
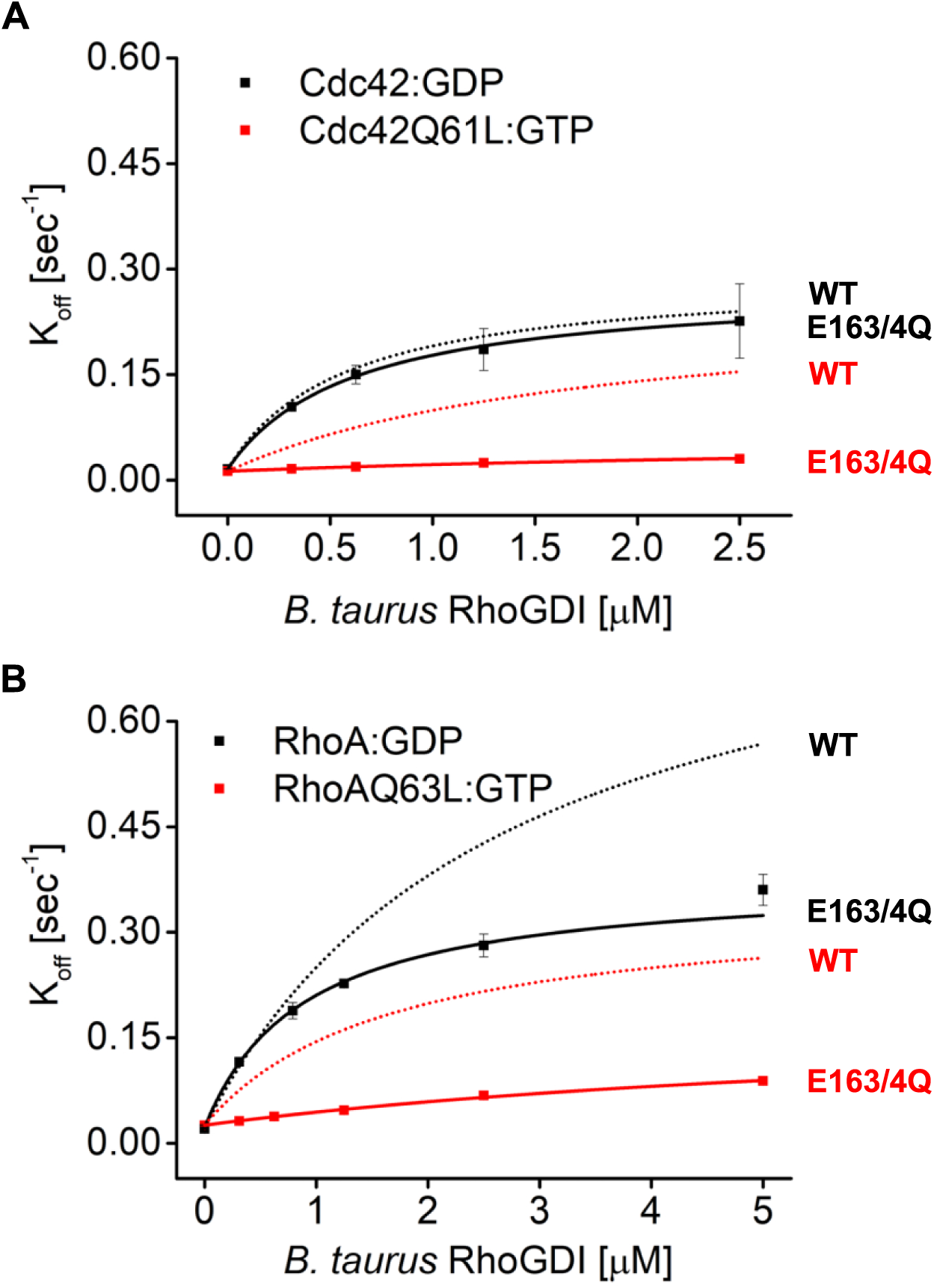
Mutant E163/4Q bovine RhoGDI is deficient in extraction of active Cdc42 and RhoA *in vitro*. Comparison of K_off_ values obtained for inactive (Cdc42:GDP, RhoA:GDP) and constitutively-active (Cdc42Q61L:GTP, Cdc42Q63L:GTP) Cdc42 and RhoA from wash off experiments in presence of either WT (black) or E163/4Q (QQ) (red) bovine RhoGDI. Extraction rates were fitted with a hyperbolic function.

**Supplemental Figure 9:**
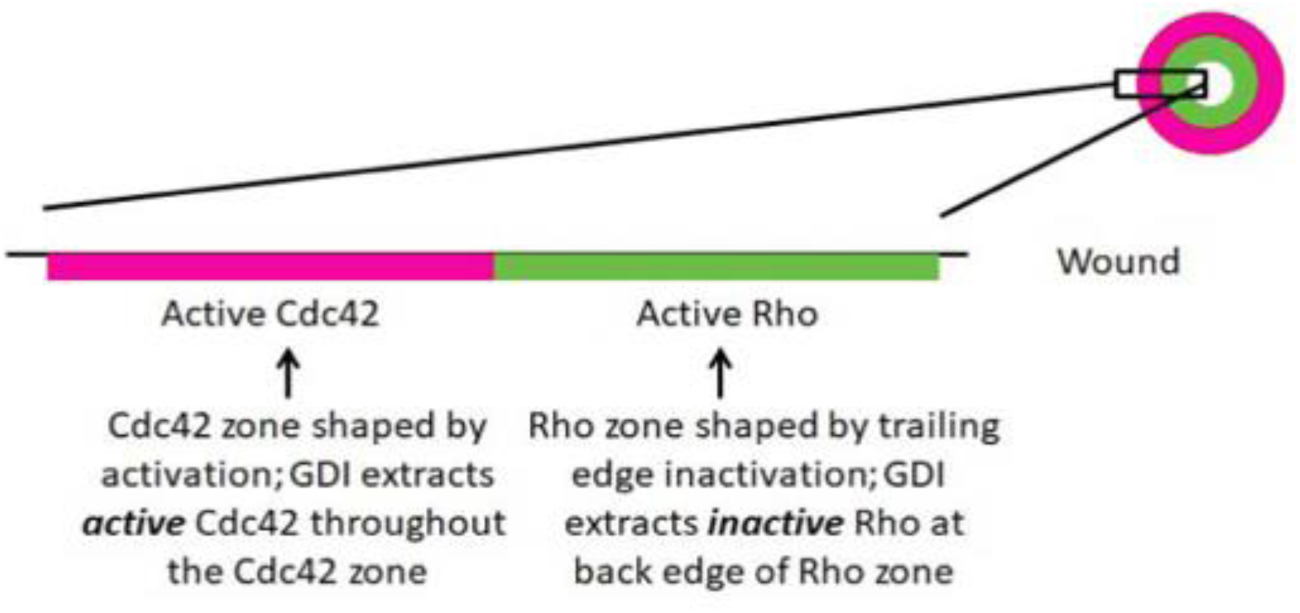
Schematic of RhoGDI’s role in RhoGTPase zone definition around wounds. Active Rho (green) and active Cdc42 (red) are activated in discrete, concentric zones that close inward as the wound heals. Cdc42 inactivation is variable throughout its zone, thus its zone is shaped by activation (Burkel et al., 2012). Conversely, the Rho zone is shaped by inactivation as it is subject to RhoGAPs 1/8 at the trailing edge of its zone (Davenport, 2016). Thus, we hypothesize that GDI extracts active Cdc42 throughout the Cdc42 zone and inactive Rho from the trialing edge of the Rho zone.

**Supplemental Figure 10:**
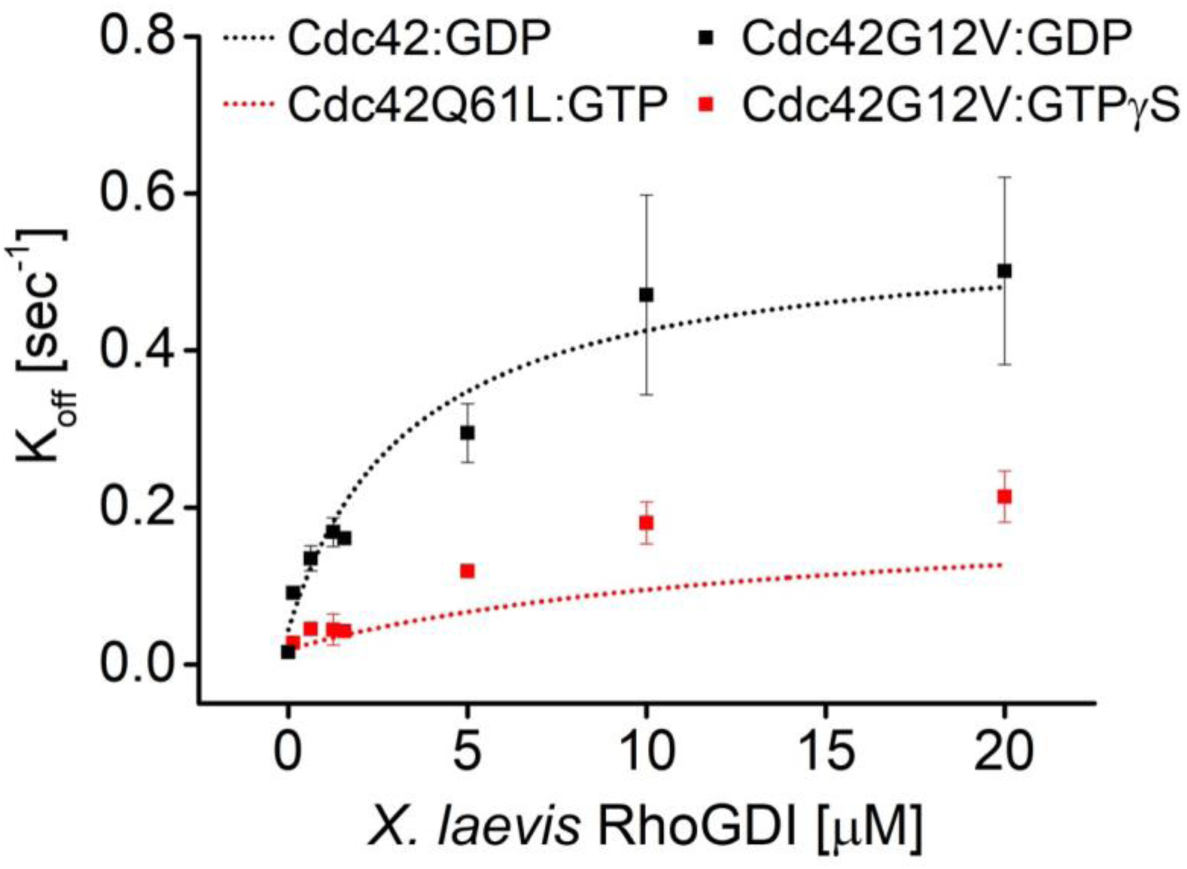
Comparison between G12V and Q61L constitutively-active RhoGTPases. K_off_ values obtained for constitutively-active Cdc42 G12V fitting the decay curves with a biexponential decay function are plotted against RhoGDI concentration. The fast rate (black) corresponds to the GDP-bound state (dotted black line), the slow rate (red) to the active GTP-state as inferred from the Q61L mutant (dotted red line).

**Supplemental Figure 11:**
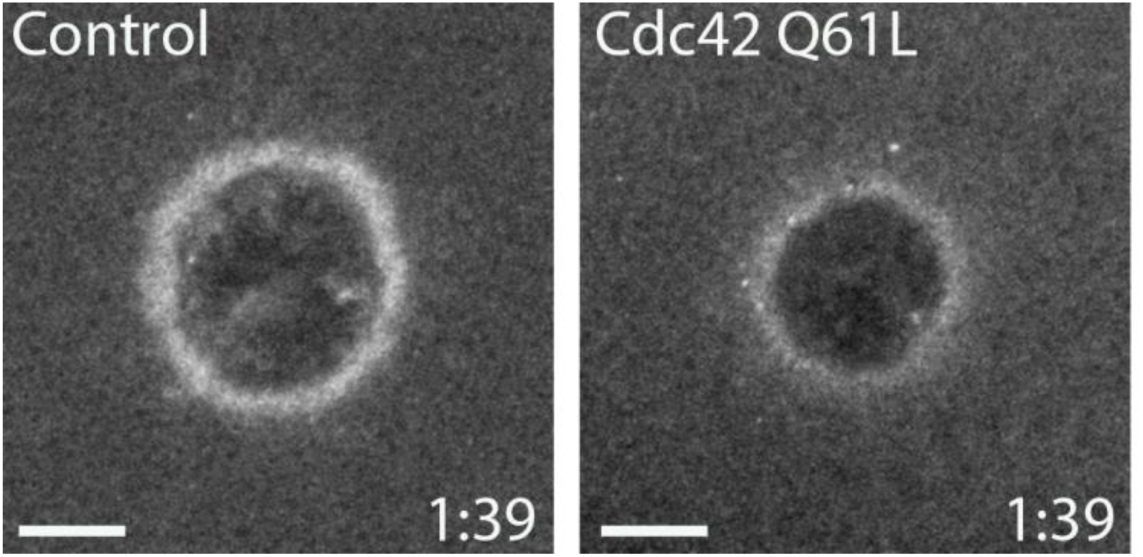
Cdc42 Q61L does not behave like constitutively-active Cdc42 *in vivo*. Oocytes injected with wGBD alone or with Cdc42 Q61L. Scale bar 10μm, time min:sec.

**Supplemental Table 1:**
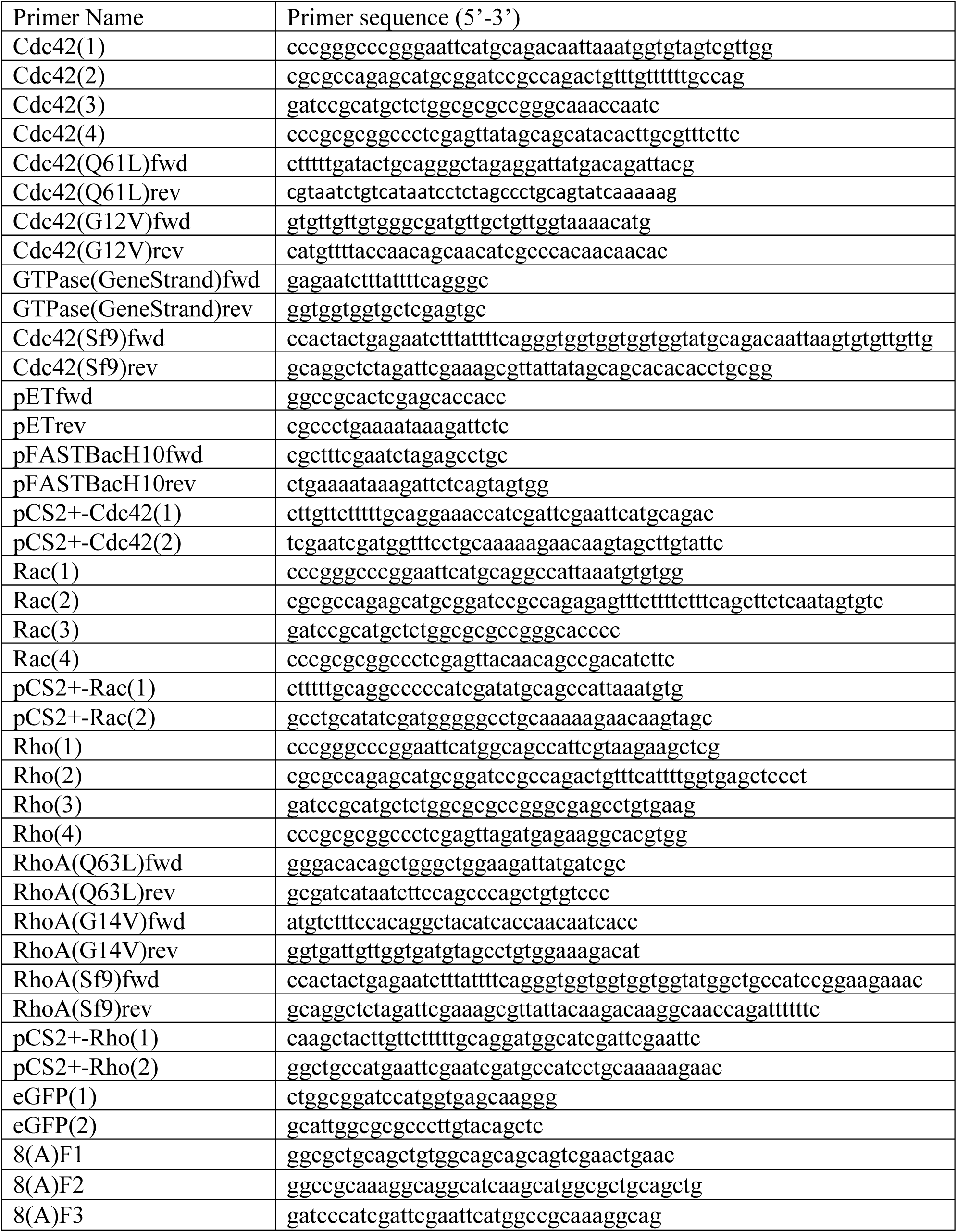
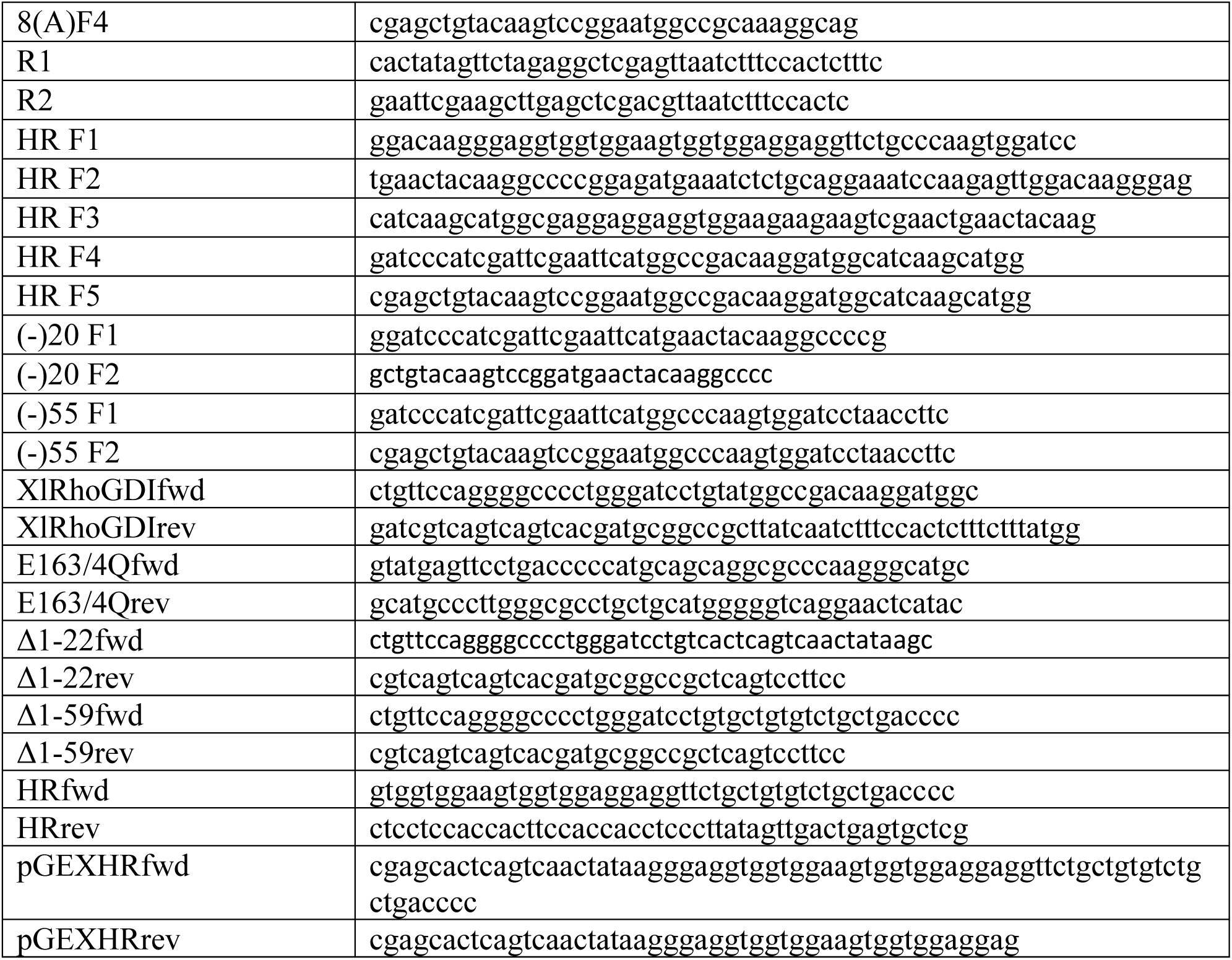
Primers.

## Supplemental Discussion

Two naturally occurring, oncogenic mutations originally discovered in Ras (Q61L and G12V) are commonly used as constitutively-active variants of Ras-like GTPases such as Cdc42 and RhoA. While often used interchangeably, it is important to note that the biochemical properties of these two single site mutations are not equivalent (Smith, Neel, & Ikura, 2013). The Q61L substitution directly affects a catalytic residue within the RhoGTPase active site and therefore directly and strongly impairs spontaneous nucleotide hydrolysis. This leads to the Q61L mutant being dominantly GTP-bound in the absence of external regulators such as GEFs and GAPs. In other words, GTPases affected by the Q61L substitution do not require GEF activity to adopt an active, GTP-bound state.

The G12V mutation, on the other hand, targets an auxiliary site important for GAP-mediated hydrolysis. GTPases affected by the G12V mutation are still capable of hydrolyzing GTP intrinsically and can thus accumulate in the inactive, GDP-bound state in the absence of external regulators. Their “constitutive activity” rather originates from a complete deficiency in inactivation via GAP-induced GTP hydrolysis. In line with this, we found that these two mutants, when purified recombinantly, accumulate in different nucleotide states as determined by HPLC analysis: Q61L mutant proteins were GTP-bound, whereas G12V ones were GDP bound after purification. We also found that the two produced different effects *in vivo* with G12V behaving as expected for a constitutively-active (ie its expression resulted in an excess of Cdc42 activity around the wound) while Q61L does not (ie its expression modestly or significantly reduced Cdc42 activity around wounds; SuppFig11).

With respect to the results obtained with these mutants *in vitro*, Q61L mutant GTPases (constitutively bound to GTP) showed monophasic membrane dissociation kinetics upon addition of GDI with rates that were clearly slower than those obtained for inactive GTPases (Fig7). This shows that the Q61L mutant uniformly adopts an active, GTP-like state distinct from the GDP form.

Since the G12V mutant was bound to GDP after purification, we first exchanged its nucleotide to GTPγS. GTPases prepared in this manner were however extracted by RhoGDI with biphasic kinetics with two characteristic rates, indicating the presence of two biochemically distinct GTPase species. Interestingly, the fast rate corresponded to the GDP-bound state, whereas the slow rate was nearly identical to the active GTP-state as inferred from the Q61L mutant (SuppFig10). From these observations, we drew the following conclusions: i) the G12V mutant likely hydrolyzes a considerable fraction of their associated GTPγS during the long time needed to prepare our extraction assays (often multiple hours, see Methods), ii) the “active states” adopted by Q61L (GTP, complete) and G12V (GTPγS, partial) are identical concerning GDI-mediated extraction from membranes. Because of these two reasons combined, we chose to work with the Q61L mutants as a proxy for the active GTPase state for all our *in vitro* extraction assays.

